# TUDCA modulates drug bioavailability to regulate resistance to acute ER stress in *Saccharomyces cerevisiae*

**DOI:** 10.1101/2024.11.14.623614

**Authors:** Sarah R. Chadwick, Samuel Stack-Couture, Matthew D. Berg, Sonja Di Gregorio, Bryan Lung, Julie Genereaux, Robyn D. Moir, Christopher J. Brandl, Ian M. Willis, Erik L. Snapp, Patrick Lajoie

**Author notes:** Corresponding authors: Patrick Lajoie, PhD, Department of Anatomy and Cell Biology The University of Western Ontario London, Ontario N6A 5C1 Canada; Erik L. Snapp, PhD, Janelia Research Campus of the Howard Hughes Medical Institute Ashburn, VA 20147 USA. These authors contributed equally to this work (Sarah R. Chadwick, Samuel Stack-Couture). Department of Genome Sciences, University of Washington, Seattle, USA.

## Abstract

Cells counter accumulation of misfolded secretory proteins in the endoplasmic reticulum (ER) through activation of the Unfolded Protein Response (UPR). Small molecules termed chemical chaperones can promote protein folding to alleviate ER stress. The bile acid tauroursodeoxycholic acid (TUDCA), has been described as a chemical chaperone. While promising in models of protein folding diseases, TUDCA’s mechanism of action remains unclear. Here, we found TUDCA can rescue growth of yeast treated with the ER stressor tunicamycin (Tm), even in the absence of a functional UPR. In contrast, TUDCA failed to rescue growth on other ER stressors. Nor could TUDCA attenuate chronic UPR associated with specific gene deletions or over-expression of a misfolded mutant secretory protein. Neither pretreatment with or delayed addition of TUDCA conferred protection against Tm. Importantly, attenuation of Tm-induced toxicity required TUDCA’s critical micelle forming concentration, suggesting a mechanism where TUDCA directly sequesters drugs. Indeed, in several assays, TUDCA treated cells closely resembled cells treated with lower doses of Tm. In addition, we found TUDCA can inhibit dyes from labeling intracellular compartments. Thus, our study challenges the model of TUDCA as a chemical chaperone and suggests that TUDCA decreases drug bioavailability, allowing cells to adapt to ER stress.

## INTRODUCTION

The accumulation of unfolded or misfolded proteins in the endoplasmic reticulum (ER) results in ER stress that activates several response pathways to restore homeostasis of the secretory protein folding environment. In particular, the Unfolded Protein Response (UPR) responds to both accumulation of misfolded secretory proteins in the ER lumen, as well as ER membrane lipid perturbations (Gardner and Walter, 2011; Lajoie *et al*., 2012; Volmer *et al*., 2013; Volmer and Ron, 2015; Halbleib *et al*., 2017; Karagöz *et al*., 2017; Fun and Thibault, 2020; Ho *et al*., 2020; Ishiwata-Kimata *et al*., 2021; Celik *et al*., 2023). UPR sensors activate a transcriptional response to adapt cells to restore homeostasis (Harding *et al*., 2000; Travers *et al*., 2000; Walter and Ron, 2011).

ER stress is a feature of numerous conditions, including neurodegenerative diseases (e.g. Alzheimer’s and Parkinson’s Diseases), chronic conditions (e.g. diabetes and inflammation), as well as the aging process (Jiang *et al*., 2016; García-González *et al*., 2018; Chadwick and Lajoie, 2019; Taylor and Hetz, 2020; Ren *et al*., 2021). Chronic unresolved ER stress can lead to cell death (Chawla *et al*., 2011; Rubio *et al*., 2011; Hetz and Papa, 2018; Rai *et al*., 2024) and exacerbate disease pathology. Modulation of ER stress has been proposed as a strategy to reduce the severity of associated diseases. Several pharmacologic compounds have been developed that selectively up and downregulate arms of the UPR (Maly and Papa, 2014; Gallagher and Walter, 2016; Gonzalez-Teuber *et al*., 2019; Hetz *et al*., 2019a; Marciniak *et al*., 2022). In a parallel approach, ER stress could (theoretically) be ameliorated by directly decreasing the misfolded secretory protein burden either by refolding, degrading, or decreasing the synthesis of misfolded/unfolded proteins (Hetz *et al*., 2019b; Grandjean and Wiseman, 2020; Kelly, 2020). A broad group of small molecules appear to improve the secretory protein folding environment by possibly directly assisting protein folding, stability, or trafficking. These compounds have been termed “chemical chaperones” (Papp and Csermely; Welch and Randell Brown, 1996). Importantly, the different reported chemical chaperones appear to have distinct impacts on protein folding capacity and stress coping mechanisms (Uppala *et al*., 2017).

Tauroursodeoxycholic acid (TUDCA) has been described as a chemical chaperone and shows promise in therapeutically decreasing ER stress. TUDCA is a taurine-conjugated bile acid produced in small amounts in the human body by intestinal bacteria (Winston and Theriot, 2020). It is the primary bile acid produced in Asian and North American black bears (Wang and Carey, 2014; Li *et al*., 2016). TUDCA (and bear bile acid in general) has been used for centuries as a traditional Chinese remedy (Qiao *et al*., 2011) and can improve symptoms and slow progression of numerous ER stress-associated diseases including neurodegeneration, cardiac dysfunction, retinal degeneration, and type 2 diabetes (Keene *et al*., 2002; Ozcan *et al*., 2006; Rivard *et al*., 2007; Boatright *et al*., 2009; Berger and Haller, 2011; Nunes *et al*., 2012; Lawson *et al*., 2016; Lojpur *et al*., 2019). TUDCA reportedly improves secretory protein folding diseases and attenuates UPR activation (Xie *et al*., 2002; Ozcan *et al*., 2006; Cao *et al*., 2013; Uppala *et al*., 2017). TUDCA has been hypothesized to decrease UPR activation by stabilizing protein conformation, thereby improving the folding capacity of the ER and decreasing ER stress (Ozcan *et al*., 2006; Omura *et al*., 2013). Due to the interest in TUDCA as a treatment for multiple human diseases, determining the mechanism of how TUDCA might regulate ER stress could identify the best targets for TUDCA, as well as suggesting routes to modify or enhance TUDCA activity (Kusaczuk, 2019).

To investigate the protective mechanism of TUDCA at the cellular level, we leveraged the genetically tractable budding yeast *Saccharomyces cerevisiae*. The UPR is evolutionarily ancient. While metazoans have a more complex response with five stress UPR sensors (IRE1ɑ and **β**, PERK and ATF6ɑ and **β**) (Walter and Ron, 2011; Wu *et al*., 2014), yeast have only the highly conserved Ire1 UPR sensor (Walter and Ron, 2011; Wu *et al*., 2014). In yeast, Ire1 cleaves the mRNA of the XBP1 homologue *HAC1*, which is then spliced into a shorter form that stimulates transcription of ∼400 genes involved in secretory protein folding, degradation, trafficking, and lipid synthesis (Cox *et al*., 1993; Cox and Walter, 1996; Shamu and Walter, 1996; Travers *et al*., 2000). Previous studies of proposed chemical chaperones (such as 4-PBA, DMSO, TMAO, and glycerol) have employed yeast models to assess the impact of these drugs on protein folding and trafficking, as well as downstream effects on conserved protein quality control machinery and pathways (Singh *et al*., 2007; Mai *et al*., 2018, 2019). However, several of these compounds are now considered proteostasis modulators that can either readjust protein folding and trafficking efficiency and/or directly regulate proteostasis by acting on specific stress response pathways such as the UPR (Balch *et al*., 2008; Ma *et al*., 2017). For example, treatment with 4-PBA can downregulate the UPR induced by tunicamycin in yeast (Ho *et al*., 2020). Specifically, 4-PBA increases misfolded secretory protein sorting into COPII vesicles and thus decreases levels of misfolded proteins in the ER lumen (Ma *et al*., 2017). By using yeast genetics and cell imaging assays, we sought to determine the mechanism by which TUDCA could resolve different forms of ER stress. Our study reveals a surprising mode of action for TUDCA and suggests TUDCA should no longer be classified as a chemical chaperone.

## RESULTS

### TUDCA restores growth in presence of Tunicamycin

We set out to systematically define the mode(s) of action by which TUDCA could decrease ER stress. We began by considering the possibilities that TUDCA could act by enhancing the intensity of the ER stress response, accelerating resolution of the stress response, decreasing the burden of unfolded secretory proteins, decreasing stressor levels in the cell, and/or improving cell function to counterbalance the effects of misfolded protein stress. Before dissecting mechanisms, we began with a simple readout assay. ER stress can slow or inhibit cell growth and ultimately kill cells. Therefore, we asked whether TUDCA could improve yeast cell growth in the face of misfolded secretory protein stress. Yeast cells were spotted on agar plates containing the classic ER stressor tunicamycin (Tm) (an inhibitor of secretory protein N-glycosylation) (Kuo and Lampen, 1974). N-glycosylation can increase solubility of hydrophobic domains of proteins, enable binding of ER lectin chaperones, and ensure proper trafficking along the secretory pathway (Drickamer, 1988; Varki, 2017). Inhibition of N-glycosylation results in UPR activation (Rose *et al*., 1989; Cox and Walter, 1996). We tested a range of TUDCA concentrations for their ability to rescue Tm-induced growth defects of wild-type cells. Only 5 mM TUDCA restored growth (**Figure 1A**). Unless otherwise indicated, 5 mM TUDCA was used for subsequent experiments. If TUDCA improves protein folding, we reasoned TUDCA should decrease UPR activation upon Tm treatment. To measure UPR activation, cells expressing the fluorescent UPR reporter, UPR-mCherry (Merksamer *et al*., 2008), were treated with 2.5 µg/ml Tm +/− TUDCA and assayed for UPR-mCherry expression over time (30-120 min) by flow cytometry (**Figure 1B**). As expected, UPR reporter expression increased with Tm treatment. However, at early times, cells co-treated with 5 mM TUDCA showed significantly lower UPR activation compared to cells treated with Tm alone. No significant difference in intensity was observed when the fluorescent protein reporter was expressed from a control UPR-independent GPD promoter (**Figure 1C**). Together, these results suggest TUDCA might improve growth by decreasing misfolded protein accumulation or otherwise improving the ER folding environment during Tm treatment.

**Figure 1:**
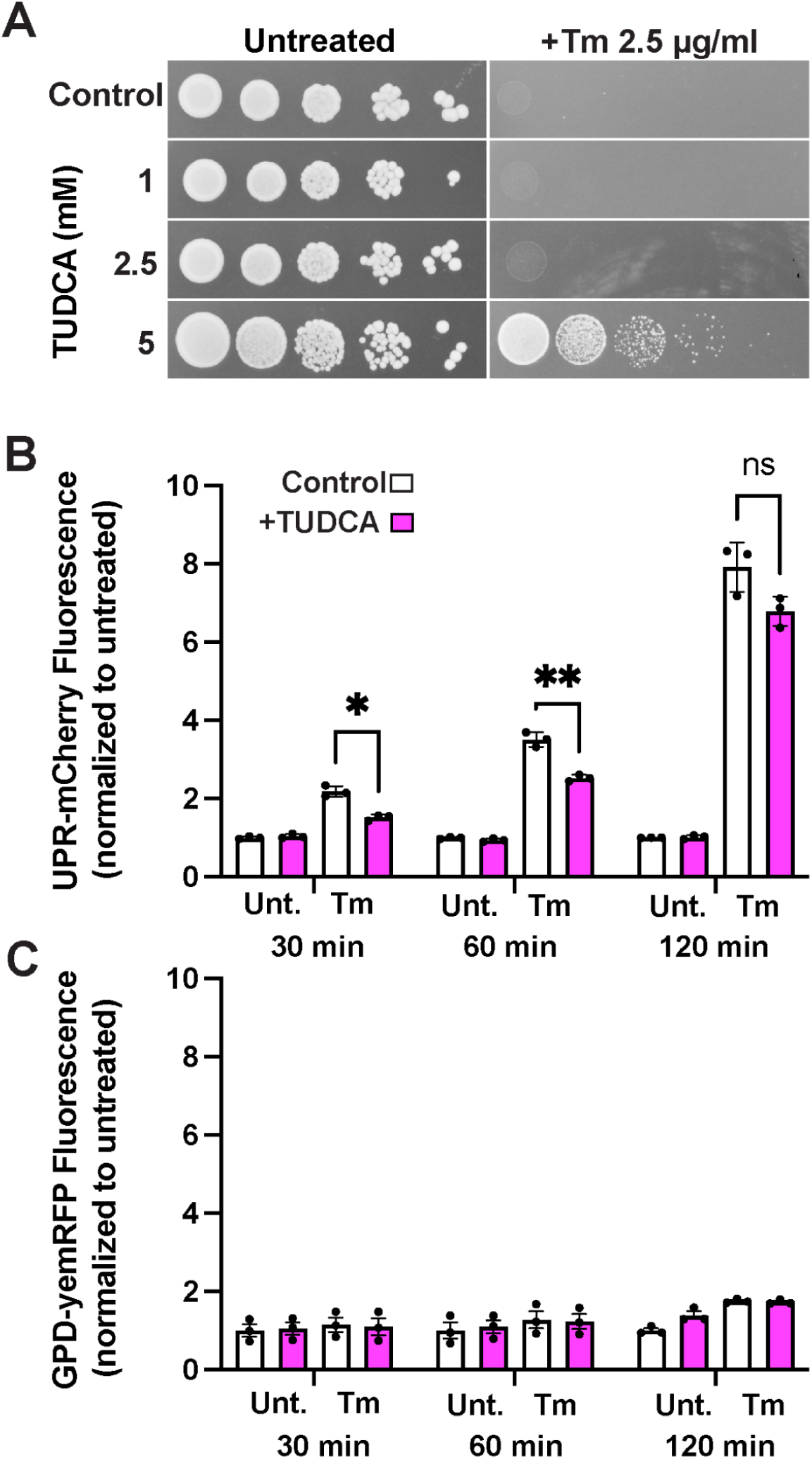
TUDCA increases resistance to Tm-induced ER stress independently of activation of the UPR. **A)** Wild-type cells were spotted on YPD plates containing 2.5 µg/mL Tm, +/− indicated concentrations of TUDCA. **B)** Wild-type cells expressing UPR-mCherry were grown to mid-log phase, and treated with 2.5 µg/ml Tm +/− 5 mM TUDCA for the indicated time periods. **C)** As a control for fluorescent protein expression, the same experiment as in (B) was conducted simultaneously with cells expressing the constitutively expressed GPD-yemRFP reporter. *p<0.05 **p<0.01 (Anova followed by Tukey’s multi comparison test)

### TUDCA alleviates phenotypes associated with acute Tm treatment

Next, we investigated TUDCA’s ability to modulate ER stress-associated phenotypes. We were interested in how broadly TUDCA rescues cells from a range of consequences of ER stress. For example, during acute ER stress induced by Tm or other stressors, small secretory proteins are retrotranslocated from the ER lumen to the cytoplasm, a phenomenon termed ER reflux (Igbaria *et al*., 2019; Lajoie and Snapp, 2020). Reflux can be observed in living cells by assessing the localization of a normally ER-localized GFP reporter, which relocalizes from the ER to the cytosol during misfolded protein stress (Igbaria *et al*., 2019; Lajoie and Snapp, 2020). Cells treated with Tm exhibited a reflux redistribution of fluorescence to the cytosol, but the reporter remained predominantly in the ER in cells co-treated with TUDCA (**Figure 2A**). Similarly, TUDCA treatment prevented other downstream consequences of Tm-induced ER stress, such as the attenuation of PKA activity (Pincus *et al*., 2014) (**Figure 2B**) and the relocalization of the transcription factor Sfp1 to the cytoplasm from the nucleus (Marion *et al*., 2004) (**Figure 2C and 2D**).

**Figure 2:**
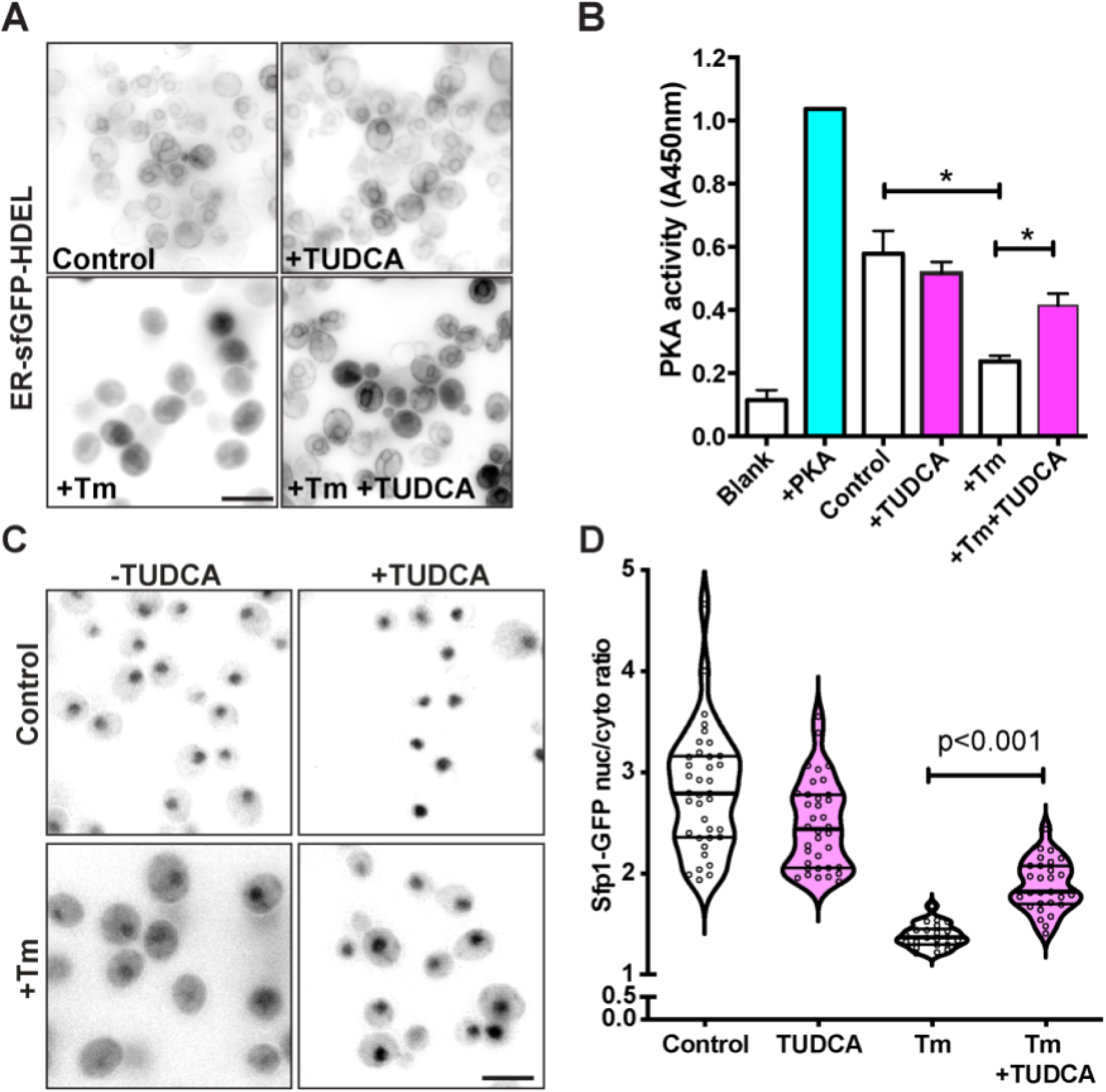
TUDCA attenuates phenotypes associated with acute ER stress. **A)** Cells expressing the fluorescent reporter ER-sfGFP-HDEL were treated with 2.5 µg/ml Tm +/− 5 mM for 2h. Bar: 10 µm **B)** Cells were treated with 2.5 µg/ml Tm +/− 5 mM TUDCA and PKA activity was measured using a colorimetric assay kit. A positive control using recombinant PKA was included. n=3 +/− SEM **C)** Wild-type cells expressing Sfp1-GFP were treated with 2.5 µg/ml Tm +/− 5 mM TUDCA and Sfp1-GFP localization in the nucleus vs. the cytoplasm was visualized using fluorescence microscopy. Bar: 10 µm. **D)** Quantitation of the Sfp1-GFP nuclear/cytoplasm ratio under the different conditions is shown in a violin plot. +/− SEM *p<0.05 (Anova followed by Tukey’s multi comparison test)

Since Tm blocks N-linked glycosylation (causing a buildup of immature glycoproteins in the ER) (Kuo and Lampen, 1974), we assessed changes in the proportion of glycosylated species of an abundant glycoprotein reporter, Pdi1 (protein disulfide isomerase 1) (Tachikawa *et al*., 1991) by immunoblot. Pdi1 has five glycosylation sites, which significantly increase its mature molecular size. Impaired glycosylation is visible with an increase in the amount of non glycosylated precursor species (lower molecular weight bands) and a decrease in the amount of the fully-glycosylated species (higher molecular weight bands). However, TUDCA did not restore glycosylation in the presence of high concentrations of Tm (1.0-2.5 µg/mL) (**Figure 3A**). Not all glycoproteins misfold if they fail to be glycosylated. To more broadly assess levels of misfolded secretory protein accumulation in the ER, we assayed changes in diffusion of the GFP-tagged Kar2 chaperone by fluorescence recovery after photobleaching (FRAP)(**Figure 3B and 3C**). The diffusion coefficient of a molecule is inversely proportional to molecular size (Einstein, 1905). Larger molecules or molecular complexes diffuse more slowly than smaller molecules. Decreased mobility of Kar2 (BiP/GRP78 in mammalian cells) directly reflects its binding to misfolded proteins (Lai *et al*., 2010; Lajoie and Snapp, 2011; Lajoie *et al*., 2012). TUDCA treatment failed to rescue low Kar2 mobility during Tm treatment. Thus, TUDCA does not prevent Tm-induced accumulation of misfolded secretory proteins.

**Figure 3:**
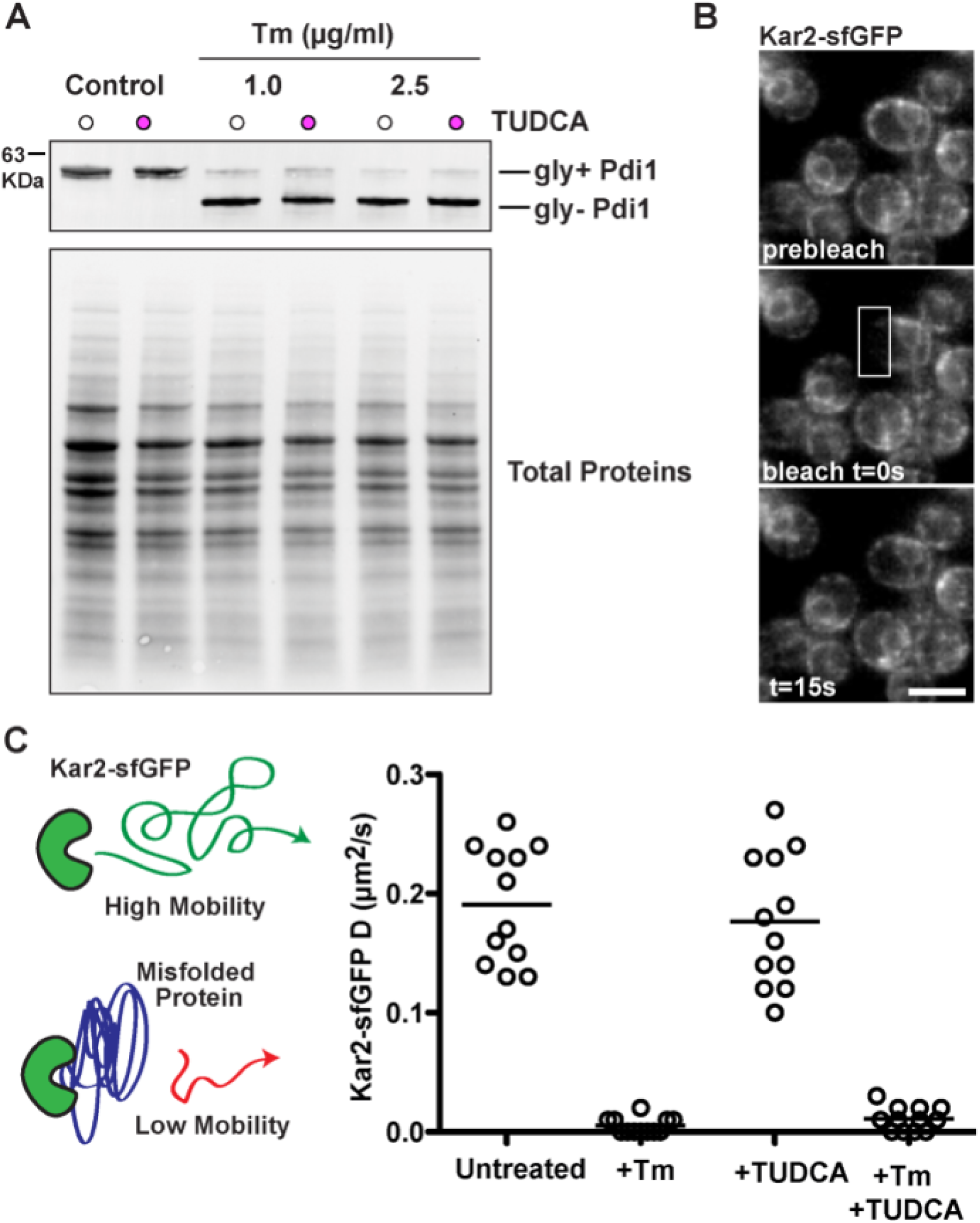
TUDCA fails to resolve N-linked glycosylation defects caused by high concentrations of Tm resulting in accumulation of misfolded proteins. **A)** Wild-type cells were grown for 2h with the indicated concentrations of Tm +/− 5 mM TUDCA. Protein was isolated and assessed by western blot for Pdi1 and total protein signal (loading control). Deglycosylated Pdi1 is seen in the lower molecular weight bands. **B)** TUDCA does not decrease the misfolded protein burden upon treatment with high concentration of Tm, as visualized by Kar2 availability measured by FRAP. Representative FRAP series of cells expressing Kar2p-sfGFP are shown. Bar: 10 μm. C) *D* values of single Kar2-sfGFP cells treated with 2.5 µg/ml Tm +/− 5mM TUDCA for 4 h are plotted on the graph.

### TUDCA differentially impact ER stressors

Impressed with the ability of TUDCA to rescue cell growth in Tm, we were curious whether TUDCA could protect against other classes of ER stressors that induce protein misfolding. Cells cannot grow in another common UPR inducer DTT, a reducing agent that prevents disulfide bond formation and acutely increases misfolded protein accumulation (Tatu *et al*., 1993). Treatment with TUDCA failed to restore growth of cells in liquid cultures treated with DTT (**Figure 4A**). We next asked whether the UPR is required for survival of cells experiencing ER stress, chronic UPR activation, even in the absence of protein misfolding, impairs cell growth (Chawla *et al*., 2011; Rubio *et al*., 2011). TUDCA did not improve, and appeared to further impair growth of cells expressing the constitutively active version (spliced) of Hac1 (*HAC1i*) (**Figure 4B**). Taken together with the results in Figure 1B, TUDCA does not appear to prevent UPR activation by another pharmacologic stressor or temper the negative consequences of UPR activation.

**Figure 4:**
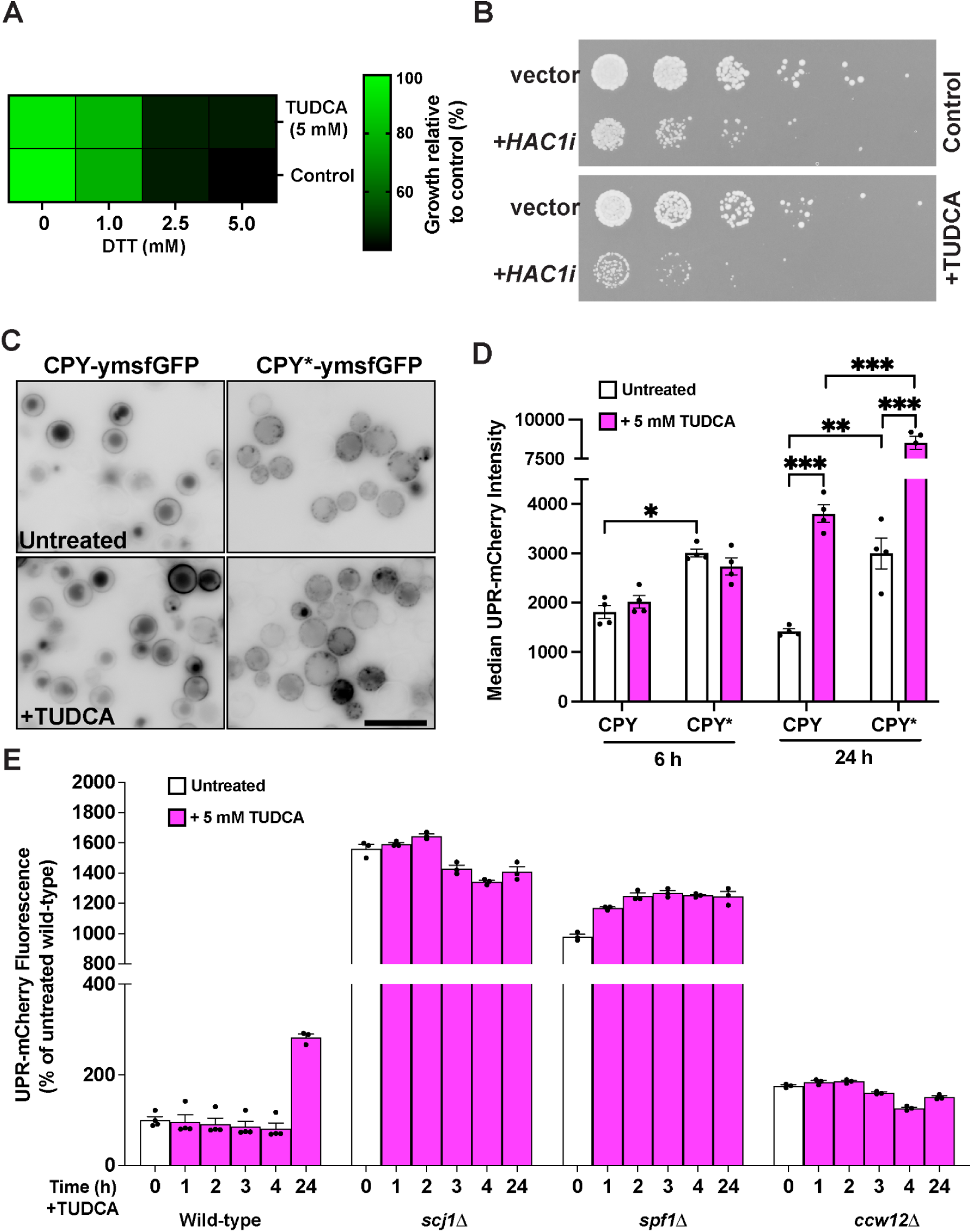
TUDCA fails to improve growth in presence of DTT or resolve genetically-induced ER stress. **A)** Wild-type cells were treated with indicated concentrations of DTT in presence or absence of 5 mM TUDCA. Growth from liquid cultures over 24 h is presented in a heat map. **B)** Cells expressing *HAC1i* or an empty vector were spotted on agar plates +/− 5 mM TUDCA. **C)** Cells expressing either galactose inducible versions of CPY or CPY* tagged with ymsfGFP were induced in galactose for 6 h +/− 5 mM TUDCA before imaging. While the native version CPY localizes to the vacuole (dark circle within a cell), the CPY* mutant is retained in the ER where it forms distinct aggregates, even in the presence of TUDCA. Bar: 10 µm **D)** Cells expressing galactose-inducible versions of untagged CPY and CPY* and UPR-mCherry were induced in galactose for 6 or 24 h +/− 5 mM TUDCA before analyzing by flow cytometry. Median UPR-mCherry fluorescence is shown in the bar graph. n = 4 +/− SEM. *p<0.05, **p<0.01, ***p<0.005. (Anova followed by Tukey multi comparison test) **E)** Wild-type and cells harboring deletions in genes causing constitutive UPR activation (*SCJ1*, *SPF1*, *CCW12*) expressing UPR-mCherry were grown to mid-log phase and treated with 5 mM TUDCA for the indicated time periods. Median fluorescence intensity of the reporters was measured by flow cytometry and mean fluorescent intensity values +/− SEM were determined for each condition (n = 3).

Next, we considered whether TUDCA could prevent or attenuate UPR induced by a single misfolded protein species. We reasoned that perhaps DTT might be too catastrophic of a stressor to correct. Therefore, we overexpressed a well characterized misfolded mutant secretory protein. A single point mutation in the vacuolar carboxypeptidase Y (CPY) referred as CPY*, leads to its misfolding, retention, aggregation within the ER, UPR activation, and subsequent degradation via ERAD (Finger *et al*., 1993; Spear and Ng, 2003). Treating cells with TUDCA for up to 24 h failed to correct CPY* defects. CPY* still did not traffic out of the ER into the vacuole (**Figure 4C**). Nor did TUDCA treatment prevent CPY*-induced UPR activation. Indeed, prolonged TUDCA treatment *exacerbated* ER stress associated with CPY* and even with wild-type CPY overexpression (**Figure 4D**). Finally, we evaluated the impact of TUDCA on an entirely different category of ER stress, the stress that arises from the deletion of specific genes (Jonikas *et al*., 2009). We asked if TUDCA treatment could attenuate stress in several deletion mutant strains with the UPR-mCherry reporter. *spf1*Δ cells (lack an ER ion transporter/ATPase) (Cronin *et al*., 2002), *scj1*Δ cells (lack an ER Hsp40 chaperone protein) (Silberstein *et al*., 1998), and *ccw12*Δ cells (lack a cell wall mannoprotein, which results in sensitivity to cell wall and osmotic stress) (Ragni *et al*., 2011) were treated with TUDCA for up to 24 h (**Figure 4E**). TUDCA did not alter UPR-mCherry expression in any of these cells. Thus, the benefits of TUDCA treatment appear to be limited.

### Identification of genes required for Tm stress mitigation by TUDCA

Together, our results led us to hypothesize that TUDCA only enabled cells to overcome stress associated with the action of Tm. Therefore, we narrowed our focus to identifying the molecular mechanism of how TUDCA rescued cell growth from the effects of Tm.

In an attempt to identify genes required for TUDCA’s action, we screened the yeast non-essential deletion library (Giaever *et al*., 2002) for deletions that modulate the rescue of Tm-induced growth defects by TUDCA (**Figure 5A**). In accordance with our results that a functional UPR is still required for TUDCA to rescue growth at high Tm concentrations (**Figure 1**), UPR genes were identified in this screen (*IRE1*, *HAC1*). Interestingly, several genes required for TUDCA rescue of Tm treatment were related to other stress responses, including *SLT2*, *MID2, BCK1* and *RLM1,* which are part of the cell wall integrity pathway and response to cell wall stress, *HOG1* and *PTC1,* which regulate the response to osmotic stress, and *CNB1* which regulates the calcineurin pathway (**Figure 5B and Supplemental File 1**). When analyzed by functional properties using TheCellMap (Costanzo *et al*., 2016; Usaj *et al*., 2017), the majority of gene deletion mutants treated with Tm, but not rescued by TUDCA were involved in glycosylation, protein folding, and the cell wall (**Figure 5C**). Thus, our screen data reveal that TUDCA does not replace the need for stress responses required to cope with Tm-induced stress, including the UPR.

**Figure 5:**
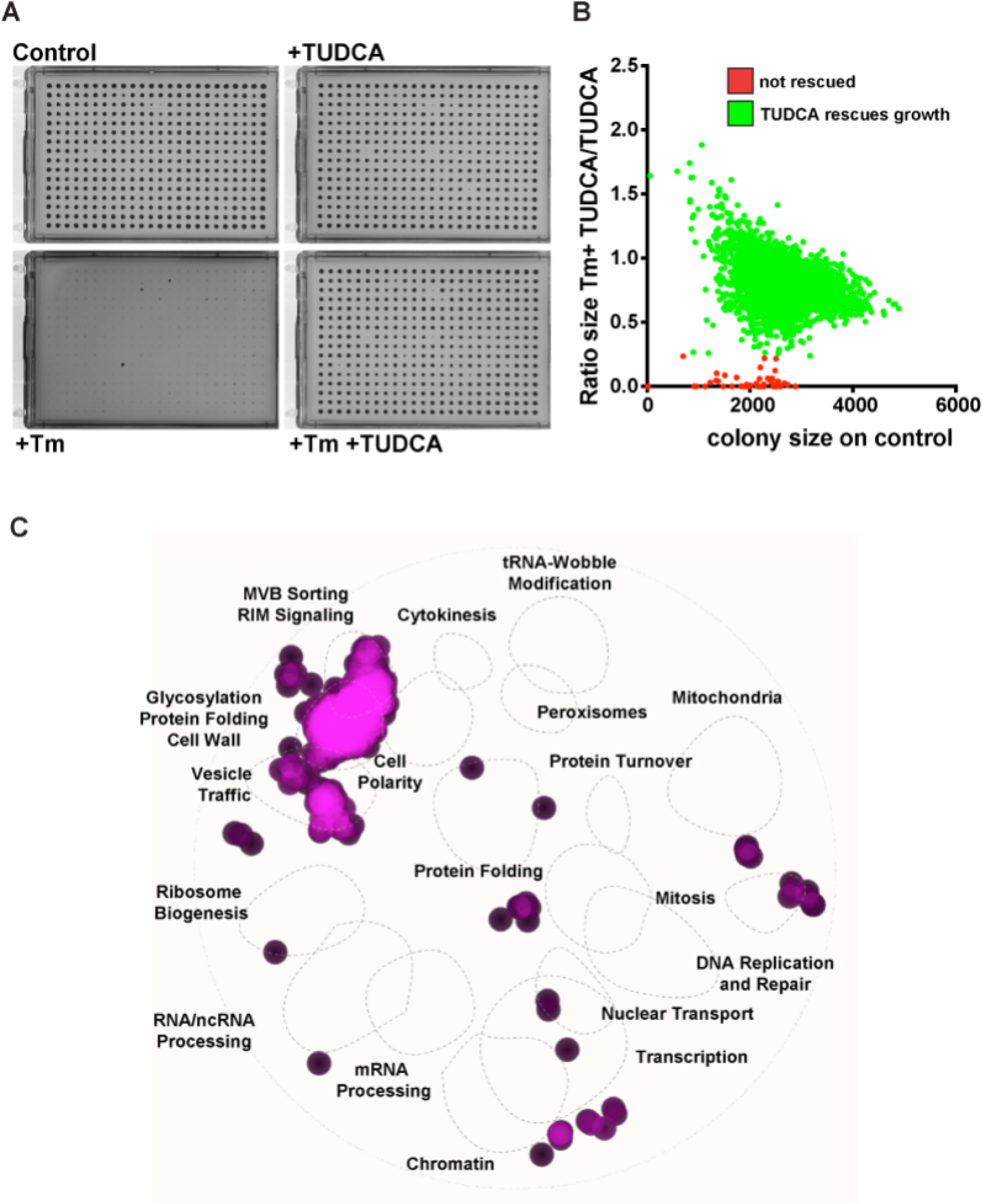
Comprehensive assessment of the genes required for TUDCA to restore growth on Tm. **A**)The yeast library of strains deleted for non-essential genes before pinning on plates containing the 2.5 µg/ml Tm and TUDCA. Colony size on the control plate was quantified and plotted against the ratio of colony sizes on the Tm plate vs. Tm + TUDCA to determine whether growth was rescued by addition of TUDCA. **B**) A colony size ratio of < 0.25 was defined as “Not rescued”, and gene deletions in this category were identified and classified (**C**) General biological processes associated with genes essential for TUDCA rescue of Tm-induced growth defect, as determined by Spatial Analysis of Functional Enrichment (SAFE).

### TUDCA mitigates Tm-induced growth defect independently of the UPR

If TUDCA does not prevent gross accumulation of unglycosylated proteins or activation of the UPR, we asked whether TUDCA might act more subtly. To explore TUDCA’s mechanism of action, we examined TUDCA’s effects during treatment with low levels of Tm. Fortunately, Tm effects are titratable over concentrations of more than an order of magnitude. Even low drug concentrations partially inhibit N-glycosylation and activate detectable UPR (Rutkowski *et al*., 2006; Lajoie *et al*., 2012). We used the Pdi1 glycosylation assay to measure Tm activity and found that even 0.1 µg/ml Tm was sufficient to inhibit N-glycosylation of the majority of Pdi1 within 2 h (**Figure 6A**). However, at these lower Tm concentrations (0.1-0.5 µg/ml), TUDCA now prevented Pdi1 deglycosylation (**Figure 6A and B**). In agreement with less accumulation of misfolded protein, we also observed less Tm-induced UPR in yeast expressing the UPR-mCherry reporter (**Figure 6C**). No change in fluorescence was observed for yeast constitutively expressing the GPD-driven yemRFP negative control (**Figure 6D**). Interestingly, these data suggest that TUDCA increases the concentration of Tm required to induce UPR. A direct prediction of this hypothesis is that TUDCA treatment could reduce the need for a functioning UPR to survive normally lethal low doses of Tm. Cells without a functional UPR (*ire1*Δ and *hac1*Δ) are especially sensitive to ER stress (Cox and Walter, 1996; Babour *et al*., 2010) and unable to grow on Tm plates, even at low concentrations. Indeed, we found that TUDCA could lower Tm toxicity sufficiently to reduce the requirement for UPR genes to enable yeast growth at a lower concentration (0.25 µg/ml) of Tm, but not two-fold higher at 0.5 µg/ml (**Figure 7**). Thus, TUDCA can improve cell resistance to Tm independently of the UPR. Together, those data suggest that at lower concentrations of Tm, TUDCA decreases the effects of Tm to allow adaptation overtime and growth.

**Figure 6:**
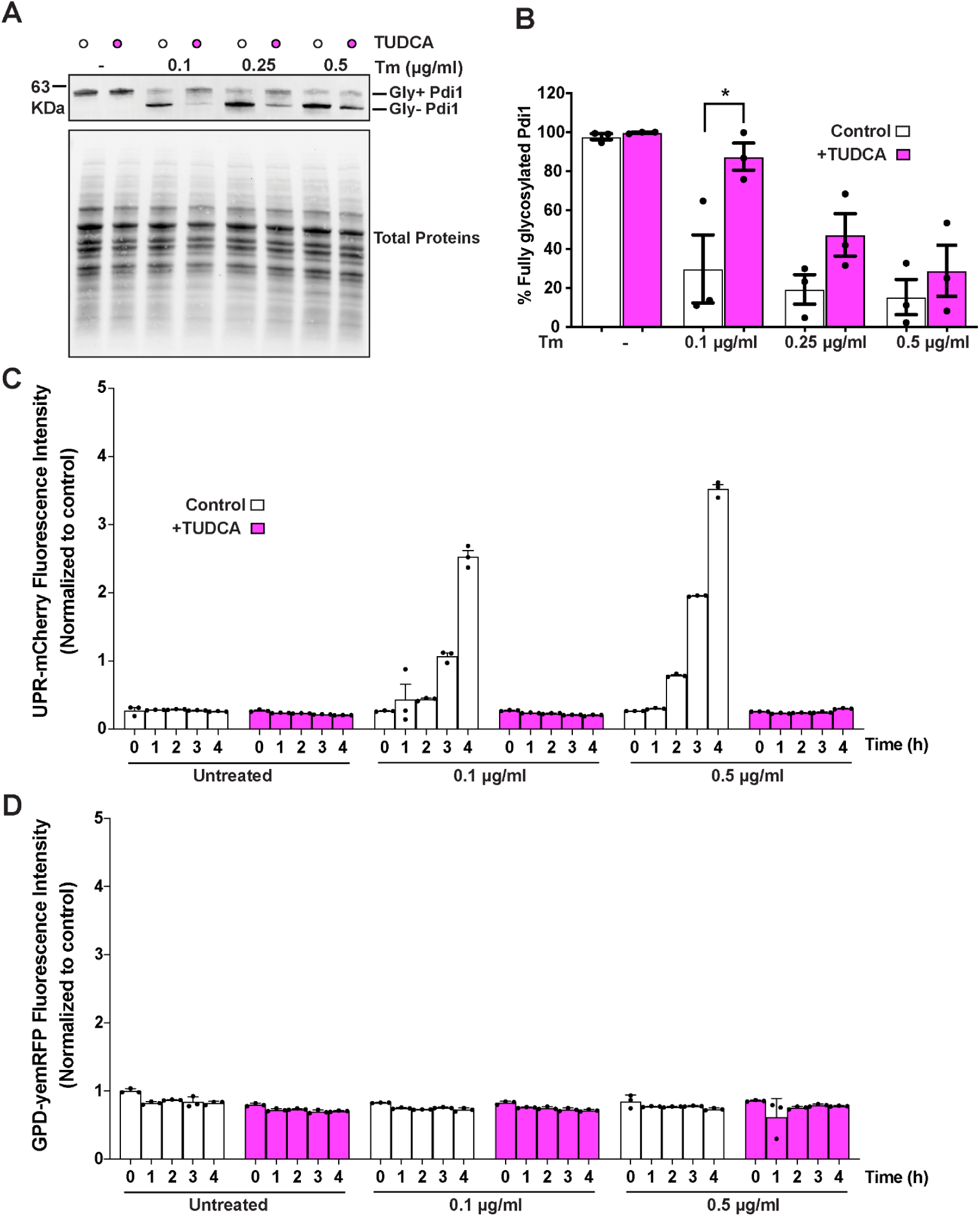
TUDCA decreases UPR activation during Tm-induced ER stress at lower concentration of stressor. **A)** TUDCA relieves N-linked glycosylation inhibition caused by low concentrations of Tm. Wild-type cells were grown for 2 h with the indicated concentrations of Tm +/− 5 mM TUDCA. Whole cell lysates were separated by SDS-PAGE and immunoblotting with anti-Pdi1 and stain-free signal (control). Unglycosylated Pdi1 is seen in the lower molecular weight bands. **B)** Densitometric analysis was performed for each Tm concentration and normalized using total protein stain. Percentage of glycosylated Pdi1 (upper bands) vs. unglycosylated Pdi1 (lower bands) was calculated for each condition and represented +/− SEM. Statistical analysis was performed using t-tests (n = 3). *p<0.05. (Anova followed by Tukey multi comparison test) **C)** Wild-type cells expressing UPR-mCherry were grown to mid-log phase and treated with indicated Tm concentrations +/− 5 mM TUDCA for the indicated time periods and fluorescence assessed by flow cytometry. Median Fluorescence intensity is shown in bar graph +/− SEM. **D)** As a control for fluorescent protein expression, the same experiment as in (C) was conducted simultaneously with cells expressing the constitutively expressed GPD-yemRFP reporter.

**Figure 7:**
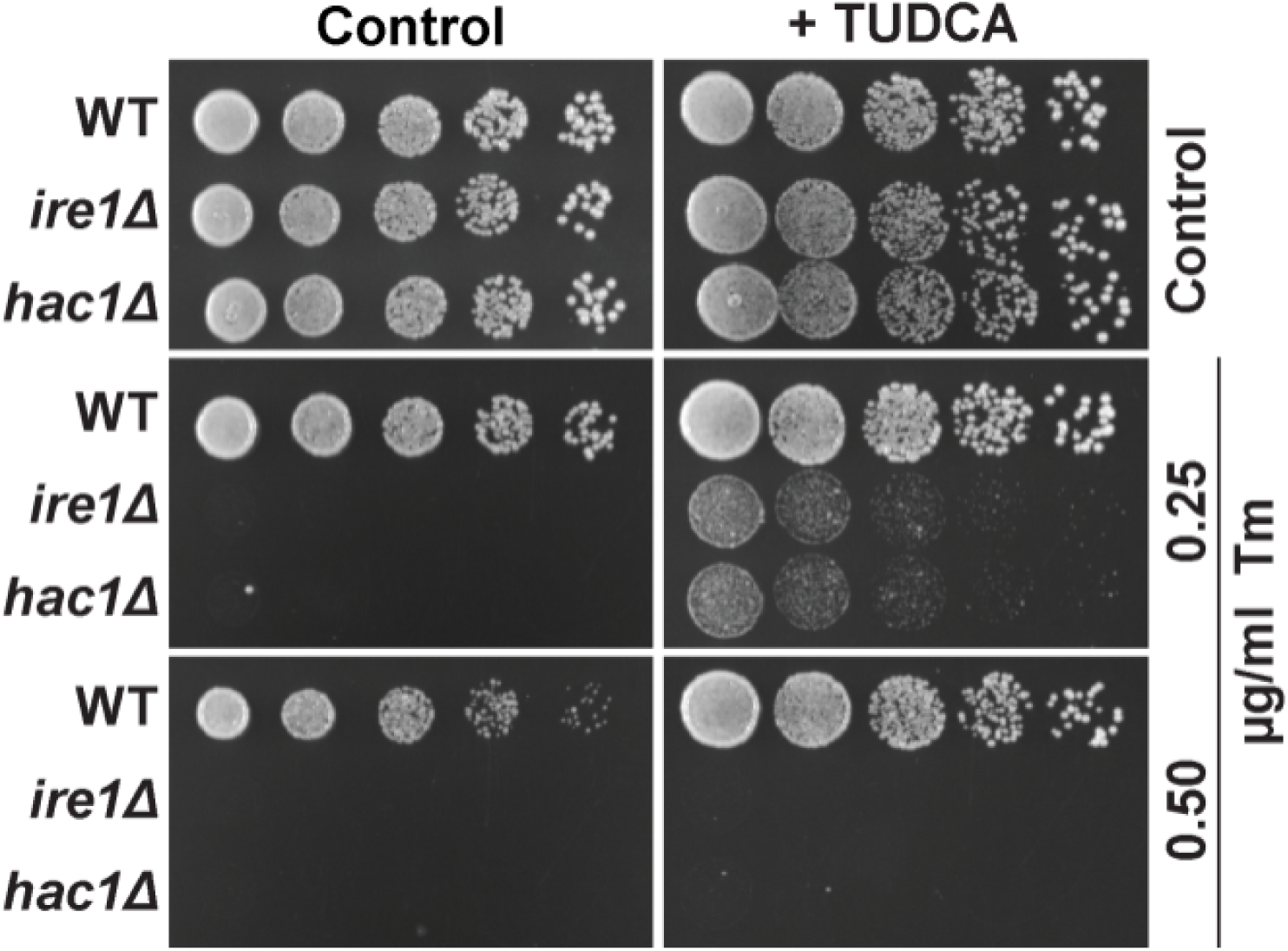
TUDCA increases resistance to Tm stress independently of the UPR. Wild-type cells and those harboring deletions of genes encoding UPR effectors *IRE1* and *HAC1* were spotted on YPD plates containing 0.25 or 0.50 µg/mL Tm +/− TUDCA.

In light of the results from Figures 6 and 7, we revisited our findings in Figure 1. We hypothesized that TUDCA might protect cells from high doses of Tm by decreasing the effective Tm dose experienced by the cells and/or by accelerating the rate of adaptation to Tm stress. To test the latter hypothesis, we treated yeast expressing the UPR-mCherry reporter with 2.5 µg/ml Tm +/− TUDCA and then measured mean reporter intensity at several time points. By 240 minutes, the TUDCA + Tm cells experienced significantly less stress than cells treated with Tm alone (**Figure 8A**). We measured the total area under the curve (**Figure 8B**) and further reinforced the interpretation that TUDCA decreased the total stress experienced. The UPR-mCherry reporter is not turned over and is a measure of total stress experienced, but does not indicate current stress level. To assess whether the rate of stress resolution changed with TUDCA treatment, we directly assayed splicing of the UPR effector, *HAC1* mRNA by activated Ire1 (**Figure 8C**). At 2 h, similar levels of *HAC1* splicing were observed in response to Tm with or without TUDCA. Yet, at 6 hours, *HAC1* splicing had decreased substantially in the presence versus the absence of TUDCA. In contrast, without TUDCA, Tm-induced *HAC1* splicing only began to dissipate after 6 hours. Thus, TUDCA decreased the total stress experienced and accelerated stress resolution.

**Figure 8:**
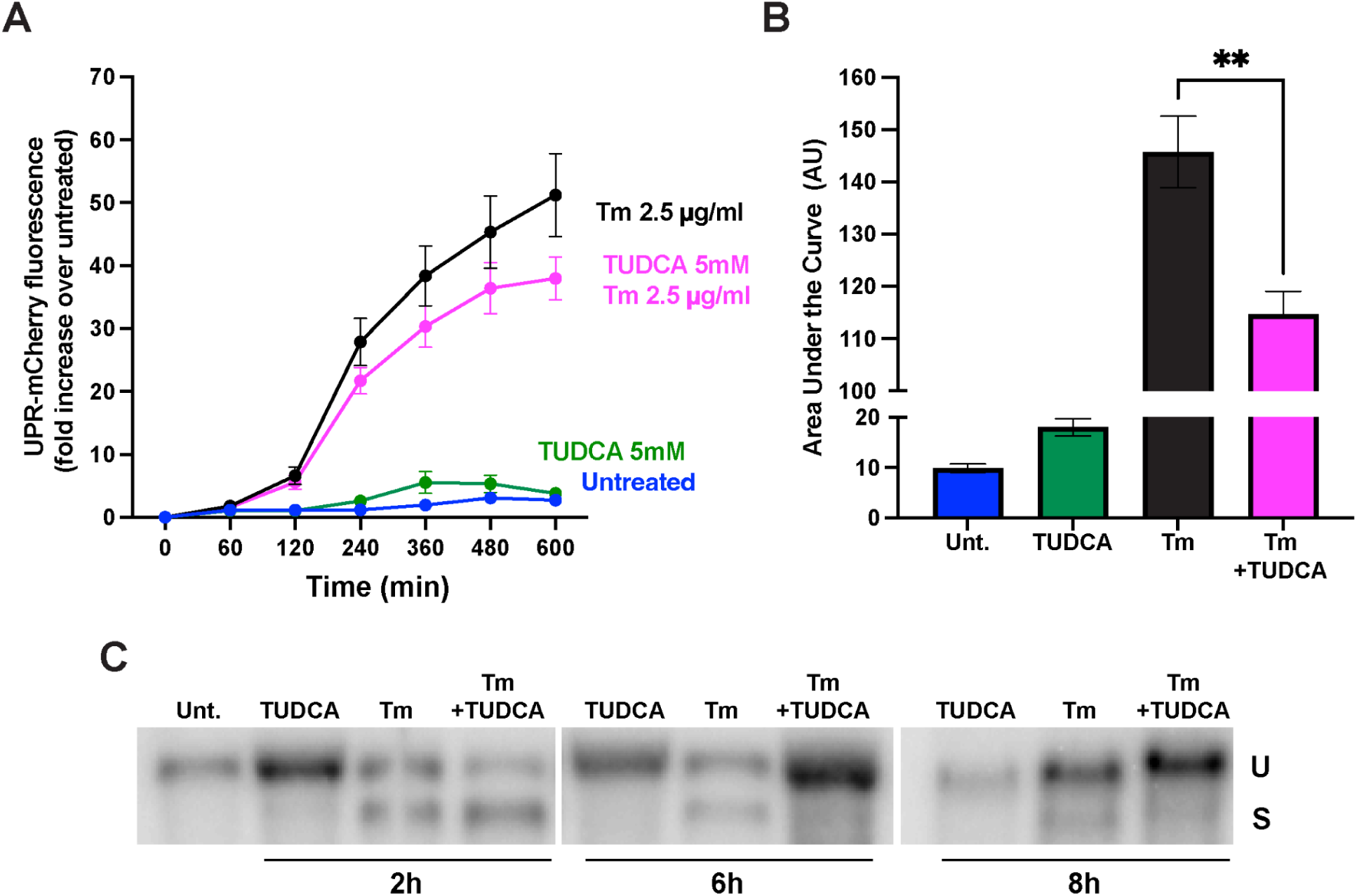
TUDCA decreases total UPR and accelerates UPR resolution during Tm-induced ER stress. **A)** Wild-type cells expressing UPR-mCherry were treated with 2.5 µg/ml Tm +/− 5 mM TUDCA for the indicated time periods and median fluorescence intensity was measured over time by flow cytometry n = 3 +/− SEM. **B)** Area under the curve is shown in the bar graph. **p<0.01. (Anova followed by Tukey multi comparison test) **C)** TUDCA accelerates attenuation of *HAC1* splicing. RNA was isolated from untreated cells or cells treated with 1.0 µg/ml Tm for 2, 6 and 8 hours and processed for Northern blot. U, unspliced *HAC1* mRNA S, spliced *HAC1* mRNA.

### Acute TUDCA treatment is not associated with profound transcriptional changes

We next considered the hypothesis that TUDCA reprograms cells to resist Tm. We complemented our genetics screen with a transcriptional profile of cells treated with TUDCA for 2 h, a time point when TUDCA treatment decreases UPR activation in cells treated with low doses of Tm (**Figure 6C**). We anticipated that a TUDCA protective stress response would be apparent by this time. Unexpectedly, TUDCA had only a modest effect on the yeast transcriptome. We identified 30 upregulated and 29 downregulated genes (adjusted *p-value* < 0.05) following TUDCA treatment (**Supplemental Figure 1 and Supplemental File 2**). Differentially expressed genes were enriched for components of the cell periphery (cell wall/plasma membrane) and for active drug transporters (**Supplemental Figure 1B**). The plasma membrane ATP-binding cassette (ABC) transporter genes (*PDR5, YOR1, SNQ2*) involved in multidrug resistance were upregulated. Interestingly, overexpression of *PDR5* has been previously reported to alleviate pharmacologically-induced ER stress in yeast (Schmidt *et al*., 2019). We hypothesized that the transporters could be upregulated by TUDCA to stimulate efflux of Tm. However, a mutant strain with all 3 transporters deleted only modestly decreased the ability of TUDCA to rescue the Tm-induced growth defect compared to the wild-type (**Supplemental Figure 2**). The difference may be unrelated to TUDCA effects as the transporter mutant strain has increased sensitivity to Tm (Rogers *et al*., 2001). Thus, the results do not support a model of TUDCA protection from Tm by multidrug transporter upregulation. Interestingly, differentially expressed genes were enriched for the GO oxidoreductase activity category; TUDCA has been previously reported to regulate oxidative stress in other organisms (Wei *et al*., 2008; Cremers *et al*., 2014; Zhang and Wang, 2018; Hou *et al*., 2021; Pioltine *et al*., 2021). Thus, we tested the ability of TUDCA to alleviate oxidative stress. It was previously reported that a conserved cysteine within the ER chaperone Kar2 plays a critical role in sensing oxidative stress, and that mutation of Kar2’s Cys63 to an alanine residue confers sensitivity to diamide (Wang *et al*., 2014; Xu *et al*., 2016). TUDCA treatment improved the growth of cells expressing wildtype *KAR2* and the *kar2 C63A* mutation in the presence of diamide (**Supplemental Figure 3A**). However, TUDCA did not rescue cells treated with hydrogen peroxide (**Supplemental Figure 3B**). Diamide causes upregulation of genes associated with the cell wall integrity pathway (Gasch *et al*., 2000) and, unlike hydrogen peroxide, induces significant changes to the cell wall (Vilella *et al*., 2005). Thus, TUDCA can have differential effects on various chemicals and ER stressors. The absence of profound transcription changes in response to TUDCA argues that it must work rather quickly upon addition of Tm. Indeed pretreatment with TUDCA did not promote growth in Tm-containing plates (**Figure 9A**). Moreover, delayed addition of TUDCA following addition of Tm for 2 h did not restore growth (**Figure 9B**). Thus, in order to protect against Tm, TUDCA needs to be present together with the stressor.

**Figure 9:**
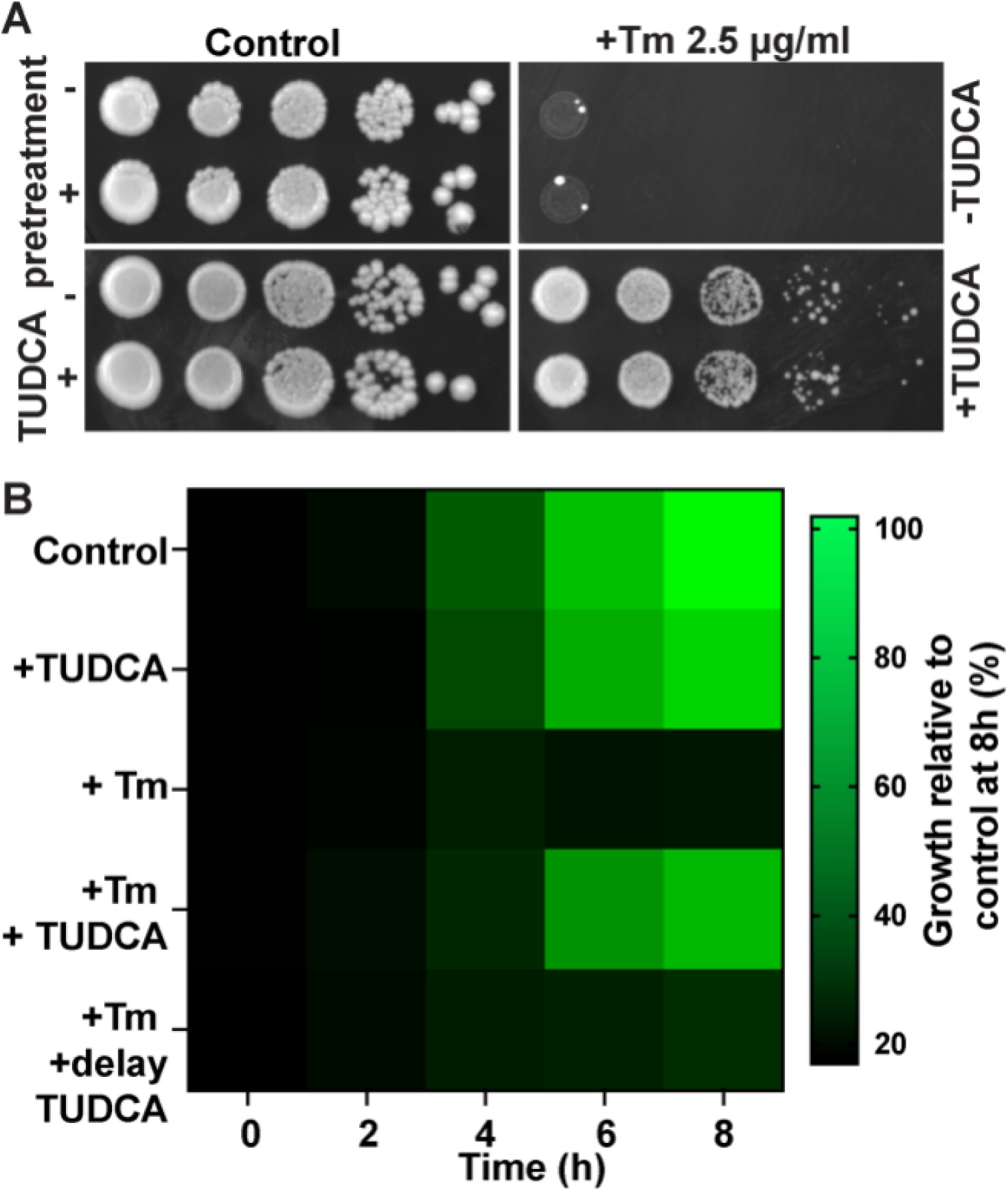
TUDCA must be added together with Tm to relieve Tm-induced growth defect. **A)** Wild-type cells untreated and treated overnight with 5 mM TUDCA were spotted on YPD plates containing 2.5 µg/mL Tm, +/− 5 mM TUDCA. **B**) Wild-type cells were grown in YPD containing 2.5 µg/mL Tm +/− 5mM TUDCA for 8 h. Alternatively, TUDCA was added 2 h (delay TUDCA) after the initial treatment with Tm. Cell growth at 8 (OD_600_) relative to control is shown in heatmap.

### TUDCA decreases drug bioavailability

Taken together, our data reveal two important aspects of TUDCA: 1) TUDCA can rescue cells from pharmacologic stressors and 2) to do so, TUDCA must be present at the time when the stressor is added. Our results suggest that TUDCA decreases intracellular drug bioavailability, probably by either modifying yeast cell wall integrity and/or directly interacting with drugs in the culture media. In support of the altered cell wall hypothesis, our transcriptome results identified cell wall components (*TIR1, TIR3, AGA1, SAG1*) among the differentially expressed genes, suggesting that TUDCA can affect cell wall composition. In support of the hypothesis that TUDCA might directly interact with Tm, other groups have reported that other bile acids can form micelles that can sequester amphiphilic drugs such as caspofungin and decrease their toxicity (Hsieh and Brock, 2017; Hsieh *et al*., 2017). Indeed, using a previously characterized assay for incorporation of coumarin-6 into micelles (Fluksman and Benny, 2019), we determined that the critical concentration for micelle formation of TUDCA is 5.17 mM (**Figure 10A**). Critically, only at micelle forming concentration were we able to observe TUDCA rescue of yeast growth in Tm (**Figure 1A**). In fact, and somewhat surprisingly, lower concentrations of TUDCA exacerbated the Tm-induced growth defect (**Figure 10B and C**) suggesting TUDCA acts differently (probably as a cell wall stressor) on cells when used below the micelle forming concentration. TUDCA is present in vast excess over Tm, as 1 µg/ml of Tm is 1.2 µM.

**Figure 10:**
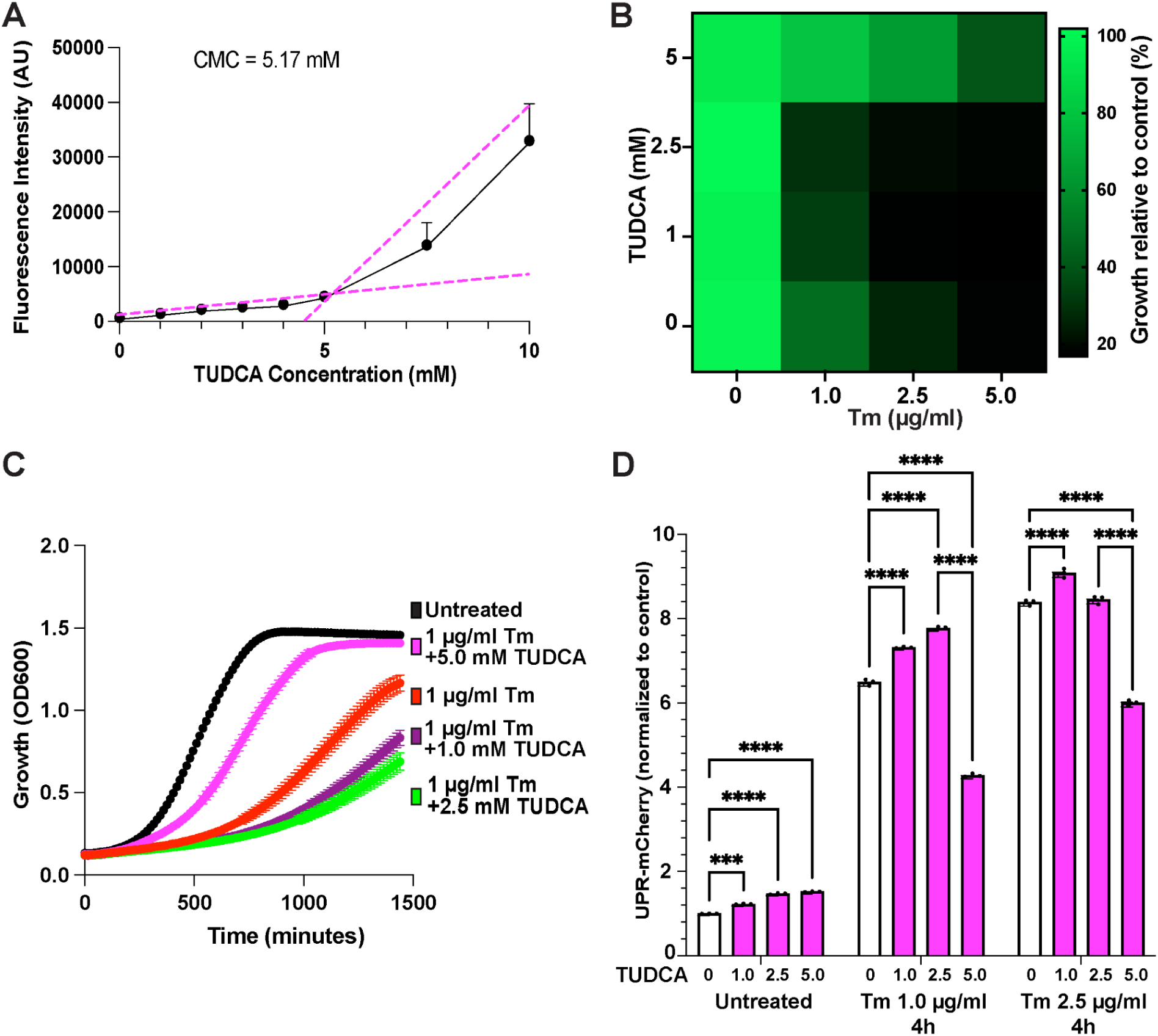
Attenuation of Tm-induced ER stress requires TUDCA’s critical micelle forming concentration. **A)** Critical micelle concentration (CMC) was determined using coumarin-6. TUDCA solutions of varying concentrations were added to 6 μM dried coumarin-6 and incubated in the dark at room temperature for 20 h. Fluorescence intensity of the samples were read at excitation and emission wavelengths of 485 ± 20 nm and 528 ± 20 nm respectively. Fluorescence intensities were plotted against corresponding TUDCA concentrations and the CMC was determined by identifying the point of intersection of the two tangents created by the graph. n=3. **B)** Wild-type cells were grown in YPD media containing different concentrations of Tm and TUDCA. Cell growth (OD_600_) was measured every 15 min for 24 h to generate growth curves. Areas under the curves are represented in a heat map for each condition. **C)** Growth curves of Wild-type cells were grown in YPD media containing different concentrations of Tm and TUDCA. **D)** Wild-type cells expressing UPR-mCherry were grown to mid-log phase and treated with indicated Tm and TUDCA concentrations for 4 h and fluorescence was assessed by flow cytometry. ***p<0.005, ****p<0.0001 (Anova followed by Tukey’s multi comparison test)

It is difficult to directly measure the intracellular content of Tm. Fortuitously, we found in the course of our studies that TUDCA also interfered with the ability of common cellular dyes to stain cells. Taking advantage of this observation, we used dye labeling as a proxy to assess TUDCA’s capacity to alter the bioavailability of compounds in general. We tested the efficacy of fluorescent dye labeling across a range of TUDCA concentrations. We found that simultaneous addition of TUDCA with either mitotracker (a mitochondrial staining dye) or FM4-64 dramatically decreased dye labeling, but *<u>only</u>* at 5 mM TUDCA, the minimal concentration that enables micelle formation (**Supplemental Figure 4**). TUDCA was present at concentrations several orders of magnitude higher than each dye (50 nM mitotracker and 8 µM FM4-64). Moreover, pretreatment with TUDCA followed by washout prior to staining was not sufficient to impair dye uptake (**Supplemental Figure 5**), suggesting TUDCA most directly impacts drug bioavailability. Together, our functional assays for Tm and visual assay for cell dyes suggest that micelle forming concentrations of TUDCA alleviates ER stress, by decreasing intracellular drug bioavailability, independently of any function of TUDCA as a chemical chaperone. Interestingly, both Tm and mitotracker are hydrophobic compounds solubilized in DMSO, while FM4-64 is hydrophilic. While it remains unclear what criteria define TUDCA susceptible compounds, hydrophobic compounds are likely to be targets, while hydrophilic compounds, such as DTT or FM4-64 are less predictable in terms of TUDCA susceptibility.

## DISCUSSION

Unmitigated ER stress is deleterious across all organisms including yeast. The UPR monitors the fitness of the ER folding environment and adjusts components of the protein quality control machinery in response to changes in the misfolded protein burden. To regulate UPR activity, several different approaches are possible. First, UPR activity can be modulated by pharmacological UPR inhibitors or activators. Several small molecules that target ER stress sensors have been developed and are currently being tested in various ER stress-associated disease models (Gonzalez-Teuber *et al*., 2019). One could also potentially employ compounds or genetic approaches that act on either the UPR or other signaling pathways that function in parallel (Valenzuela *et al*., 2018). For example, activation of the heat shock response decreases UPR activity (Liu and Chang, 2008). Alternatively, it is possible to improve secretory protein folding using chemical chaperones (Perlmutter, 2002; Rajan *et al*., 2011). The question is which class (es) of regulators does TUDCA belong to?

The ER stressor Tm inhibits Alg7, an essential enzyme required for cell viability, which encodes the major target of Tm, the dolichyl-P-dependent N-acetylglucosamine-1-P transferase in the N-glycosylation pathway. Alg7 expression is specifically upregulated in response to UPR activation (Barnes *et al*., 1984). Any compound that rescues cells from stress and impaired growth must somehow compensate for the loss of the dolichol pathway metabolites required for N-glycosylation of secretory proteins. N-glycosylation increases hydrophilicity of secretory proteins, can regulate steps in protein folding, and enables interactions with lectin partner proteins (Reily *et al*., 2019). Here, we show that TUDCA can decrease Tm-induced ER stress and improve cell growth. Surprisingly, TUDCA can do so in the absence of a functional UPR, suggesting a mechanism that circumvents the requirement for UPR signaling. Thus, TUDCA must either block Tm activity or overcome the action of Tm. This is an important finding, since few known mechanisms can overcome Tm. One such mechanism is the upregulation of *ALG7* expression. In addition, deletion or depletion of the transporter responsible for Tm uptake into mammalian cells, MFSD2A, can also decrease Tm sensitivity (Bassik and Kampmann, 2011; Reiling *et al*., 2011). Whether a similar transporter exists in yeast is unclear. Our screen did not reveal an obvious candidate. Regardless, we find TUDCA quantitatively decreases the effects of Tm inhibition of N-linked glycosylation, at least at lower doses of Tm. Our data support a model of decreased availability/uptake of Tm (and other stressors and compounds). Since disruption of multidrug efflux transporters only modestly reduced TUDCA rescue, we hypothesize TUDCA likely decreases Tm internalization. Recent analysis of Tm-resistant aneuploid mutants revealed that the mutants have increased chitin content (Beaupere *et al*., 2018). We were surprised to observe only a few changes in the transcriptome of yeast treated with TUDCA during the 2 h time frame wherein we observe Tm resistance. If TUDCA causes significant cell wall stress, we would expect changes in the expression of CWI targets, which we did not observe. Similarly, the inability of TUDCA pretreatment to protect cells from Tm suggests that TUDCA must be present with the drug and that TUDCA’s effect does not persist. These observations suggest TUDCA might affect cell wall/plasma membrane permeability via a non-transcriptional mechanism.

Interestingly, beyond the misfolded protein stress literature, there are reports of bile acids, including TUDCA, regulating drug absorption. Multiple mechanisms have been proposed to explain the ability of bile acids to modulate drug absorption, such as increasing drug solubility, regulation of drug transporters, and changing membrane permeability (Greer *et al*., 1998; Darkoh *et al*., 2010; Guan *et al*., 2011; Pavlović *et al*., 2018). While we observed upregulation of multidrug transporters following TUDCA treatment, deletion of those same transporters had little impact on TUDCA rescue of Tm treatment (**Supplemental Figure 2**). In other eukaryotic cells, TUDCA molecules can localize to the plasma membrane at the lipid water interface (Mello-Vieira *et al*., 2013; Sheps *et al*., 2021). TUDCA treatment also affects fluidity and polarity of photoreceptor membranes (Sabat *et al*., 2021) and modulates membrane permeability in liposomes (Zhou *et al*., 2009).

Alternatively, TUDCA could directly interact with drugs/stressors themselves; bile acid salts can form complexes with drugs via ion-pairing or covalent conjugation through hydroxyl and carbonyl groups, resulting in increased drug solubility and availability (Pavlović *et al*., 2018). For example, conjugation of antibiotics such as kanamycin with bile acids results in increased toxicity for *Staphylococcus aureus (Giovagnoli et al., 2017)*. Other bile acids can modulate antifungal drug toxicity by sequestering them into micelles (Hsieh *et al*., 2017). Prolonged treatment with other bile acids such as lithocholic acid can profoundly affect the yeast transcriptome, proteome and lipidome and increase longevity (Burstein *et al*., 2012; Beach *et al*., 2013, 2015). Long term treatment with bile acids can also induce aneuploidy in yeast (Ferguson and Parry, 1984) which can lead to ER stress tolerance (Beaupere *et al*., 2018). Our data (especially **Figure 10**) support a model in which TUDCA micelles decrease bioavailability of Tm or other compounds. That is, at sufficiently high levels of TUDCA, cells experience decreased concentrations of some drugs, such that no stress response is required or enabling stress adaptation or hormesis to help cells survive normally lethal drug concentrations. It will be of interest for future studies to assess the prolonged effects of TUDCA treatment on the ER proteostasis network.

The path by which we have ultimately come to understand the mechanism of TUDCA highlights the value of a broad and systematic methodology for assessing future chemical chaperone candidates. Protection from stressors can involve mechanisms that have little to do with protein folding or stress pathways. We propose employing a combination of assays that measure not just cell stress reporters and growth assays, but directly measure protein damage, misfolding, and restoration of function across a range of stressor and putative chemical chaperone concentrations and time points. In several cases, protective effects only became apparent at longer time points (**Figures 8** and **10**) and lower doses of stressor (**Figures 6** and **7**). Similarly, other assays of cell stress or misfolded protein accumulation (**Figures 2** and **3**) gave distinct readouts of cell stress or actual levels of misfolded proteins. Any single assay could lead to a misleading conclusion. Our studies do not rule out potential cell or tissue protective roles for TUDCA in disease, but our studies provide little support for a chemical chaperone mode of action by TUDCA.

## MATERIALS AND METHODS

### Drugs

H_2_O_2_ (426000010), DTT (R0861), MitotrackerRed (M46751), FM4-64 (F34653) were purchased from ThermoFisher. TUDCA (580549), Tunicamycin (11089-65-9) and Diamide (10465-78-8) were from MilliporeSigma.

### Strains, plasmids and cell culture

Strains and plasmids used in this study are listed in **Supplemental Tables 1 and 2**. For every experiment, cells were thawed from frozen stocks and grown on YPD or selective SC agar plates at 30°C for 48 h before transferring to liquid culture. Cells were inoculated in 5 mL liquid media in polystyrene snap cap tubes, then grown overnight at 30°C in a rotating drum. OD_600_ of cultured cells was measured using a spectrophotometer. Cells were diluted to a final concentration of OD_600_ 0.2, then serially diluted 5 times. %-fold dilutions were spotted onto agar plates and incubated at 30°C for 48 h before imaging.

Plasmids encoding CPY or CPY* (G255R) were constructed by cloning the CPY or CPY* sequences from CPY/CPY*-GFP plasmids (Promlek *et al*., 2011) into the SpeI/HindIII sites of P415-GAL1 (Mumberg *et al*., 1995). To generate fluorescently-tagged versions, yeast-optimized monomeric super folder GFP (ymsfGFP) (Jiang *et al*., 2017) was inserted downstream of the CPY/CPY* coding sequence using the HindIII/SalI sites of P415-GAL1. Primers are listed in **Supplemental Table 3**.

### Liquid growth assay

Cells were inoculated in 5 mL liquid media in polystyrene snap cap tubes, then grown overnight at 30°C in a rotating drum. OD_600_ of cultured cells was measured. Cells were diluted to a final concentration of OD_600_ 0.15 in a flat-bottom 96-well plate and loaded into a BioTek plate reader in triplicate. Plates were held at 30°C with constant shaking and OD_600_ measurements were taken every 15 min.

### Flow cytometry

Flow cytometry was used to measure fluorescence from fluorescent reporter-expressing strains as well as to measure fluorescent dye internalization. Experiments were performed using a BD FACS Celesta equipped with the FACS Diva software. Strains were grown in 50 mL flasks to mid-log phase (OD_600_ ∼0.5), then treated as indicated (Tm, TUDCA, or both) for indicated time periods before measurement of fluorescence using a PE-TexasRed filter. For time course experiments, samples were loaded in triplicate into 96-well plates and measured using the BD FACS Celesta HTS plate reader system; 20 µL samples were taken at each time point and fluorescence data for a maximum of 10,000 cells were recorded. Between measurements, plates were incubated at 30°C with constant shaking. Median fluorescence data for each sample were used for analysis. No gates were applied.

### Protein isolation and immunoblot

Whole cell protein was isolated using alkaline lysis (Kushnirov, 2000). Mid-log phase cells were treated as indicated, OD_600_ was measured, and an aliquot of cells equivalent of OD_600_=1.0 was taken from each sample. Cells were pelleted then resuspended in 200 µL 0.1 M NaOH and incubated at room temperature for 5 min. Cells were pelleted again, then resuspended in 50 µL of sample buffer (4x Laemmeli buffer, water, and 1 M DTT) before boiling at 100°C for 3 min. Samples were pelleted and the supernatant was transferred to new 1.5 mL tubes.

Proteins were loaded into BioRad TGX Stain-Free gels and separated by gel electrophoresis. Stain-free technology was used as a loading control by activating the gel with UV light for 5 min before transfer. Proteins were transferred to nitrocellulose membranes using a BioRad Turbo Blot system and imaged with UV light to obtain final stain-free images. Membranes were incubated at 4°C overnight with anti-Pdi1 antibody (Santa Cruz Biologicals (sc-57963), 1/5000 in 5% powdered skim milk) then probed with secondary antibodies (Licor goat anti-mouse) for 1 h at room temperature before imaging. Densitometry analysis was conducted using Image Lab software.

### Northern Blot

RNA levels were assessed using Northern blot as previously described (Lajoie *et al*., 2012). Briefly, early-log cells were treated with Tm and TUDCA and cells were pelleted and frozen in dry ice. Total RNA was isolated using the hot phenol method. RNA was run on formaldehyde gels and transferred to Nytran Plus membrane. Blots were probed with ^32^P-labeled oligonucleotide and imaged using 820 Phosphorimager (GE Healthcare).

### Deletion library screens

Using the ROTOR HDA robotic system (Singer Instruments), frozen stock yeast deletion libraries were thawed and spotted from liquid cultures onto agar plates using 96- or 384-pin pads. Plates were incubated at 30°C for 1-2 days, then re-pinned onto agar plates containing indicated treatments and incubated at 30°C once again. Plates were imaged using a Nikon camera dock. Colony size was quantified using ColonyImager (S&P Robotics), and ratios of growth rescue were calculated by dividing relative colony size on Tm + TUDCA plates by colony size on plates containing TUDCA alone. Deletions that showed 50% growth reduction on Tm in presence of TUDCA compared to wild type were retested in duplicate to confirm the phenotype.

### RNA sequencing and transcriptome analysis

RNA was isolated from two biological replicates from early log phase growth in either YPD or YPD containing 5 mM TUDCA using the MasterPure Yeast RNA purification kit (Lucigen) according to the manufacturer’s instructions. Total RNA sequencing was performed by Genewiz (South Plainfield, NJ). Stranded Illumina TruSeq cDNA libraries with poly dT enrichment were prepared from high quality total RNA (RIN > 8). Libraries were sequenced on an Illumina HiSeq, yielding between 100 and 124 million 150 bp paired end sequencing reads per sample. The raw data have been deposited in NCBI’s Gene Expression Omnibus (Edgar *et al*., 2002) and are accessible through GEO Series accession number GSE186390.

FASTQ files were analyzed with a customized bioinformatics workflow. Adapter sequences were trimmed using Trimmomatic (Bolger *et al*., 2014) and aligned to the *S. cerevisiae* S288C reference genome (assembly R64-2-1; https://www.yeastgenome.org/) using STAR (Dobin *et al*., 2013). Reads that were uniquely mapped to the reference genome were counted for each gene using featureCounts (Liao *et al*., 2014). Differential expression analysis was performed using the DESeq2 R package (Love *et al*., 2014) with a Benjamini-Hochberg adjusted *p*-value cutoff of <= 0.05. GO analysis was performed using the Saccharomyces Genome Database (SGD) (Cherry *et al*., 2012) GO Term Finder using p < 0.01.

### Micelle formation assay

Critical micelle concentration was measured using coumarin-6 (Fluksman and Benny, 2019). A 6 μM solution of coumarin-6 was prepared in dichloromethane (DCM) and 40 μL of the stock solution was distributed to Eppendorf tubes and allowed to evaporate in a fume hood for 30 min. TUDCA solutions were prepared in dH_2_O ranging from 0 to 10 mM concentrations. Each TUDCA solution was added to a tube containing dried coumarin-6 and rotated in the dark at room temperature for 20 h. After the incubation period, 200 μL of each TUDCA-coumarin-6 solution was transferred to the wells of a 96-well plate and fluorescence intensity was read using excitation and emission wavelength filters of 485 ± 20 nm and 528 ± 20 nm, respectively, on a Cytation5 (BioTek). The resulting fluorescence intensities were plotted against the corresponding TUDCA concentration in GraphPad Prism. The critical micellar concentration was determined by identifying the point of intersection of the two tangents created by the graph. Points on the graph are representative of the average of three replicates.

### FRAP

Live Kar2-sfGFP cells (Lajoie *et al*., 2012) were imaged on a Duoscan confocal microscope system (Carl Zeiss Microimaging) equipped with a 63×, numerical aperture 1.4 oil objective and a 489 nm, 100 mW diode laser with a 500 to 550 nm bandpass filter for GFP. FRAP was performed by photobleaching a region of interest at full laser power of the 489 nm line and monitoring fluorescence recovery over time. Diffusion coefficient (*D*) values were determined using an inhomogeneous diffusion simulation, as previously described (Siggia *et al*., 2000; Snapp and Lajoie, 2011).

### Fluorescence Microscopy

Cells were inoculated in 3 mL of liquid media and grown overnight at 30°C in a shaking incubator. 200 µL aliquots were separated into 1.5 mL tubes and treated with TUDCA for 15 minutes. Fluorescent dyes FM4-64, and Mitrotracker were added to the cell suspension and imaged. 2 µL of stained cells were added to a glass slide beneath a glass coverslip. Live cell fluorescence imaging was conducted on the Cytation5 (BioTek). FM4-64, Mitotracker and azole-conjugated dyes stained cells were imaged using the TexasRed filter using a 20× Plan Extended Apochromat. NA 0.8.

CPY/CPY*-ymsfGFP was imaged using a Zeiss Axiovert A1 wide-field fluorescence microscope with 63× 1.4 NA oil objective using a Texas Red filter (586 nm excitation/603 nm emission) and an AxioCam ICm1 R1 CCD camera. ImageJ was used to analyze the images (Schneider *et al*., 2012).

## ACKNOWLEDGEMENTS

PL is supported by an NSERC Discovery Grant (RGPIN-2022-05267), CIHR Project Grants (PJT 168882 and ARB 192062) and a Canadian Foundation for Innovation (CFI) John R. Evans Leader Fund Grant (65183). CJB was supported by an NSERC Discovery Grant (RGPIN-2015-04394). SRC and MDB held an Alexander Graham Bell Canada Graduate Scholarship from the NSERC. MDB is currently supported by a CIHR Postdoctoral Fellowship (193932). BL held an NSERC Undergraduate Research Award. ELS is supported by the Howard Hughes Medical Institute. Part of the RNAseq analysis was performed at Albert Einstein College of Medicine and was funded by an internal pilot project grant to ELS.

The authors thank Peter Walter (UCSF/Altos), Scott Moye-Rowley (University of Iowa), Carolyn Sevier (Cornell University) and Yukio Kimata (Nara Institute of Science and Technology) for providing yeast strains and plasmids.

---

**Supplemental Figure 1:**
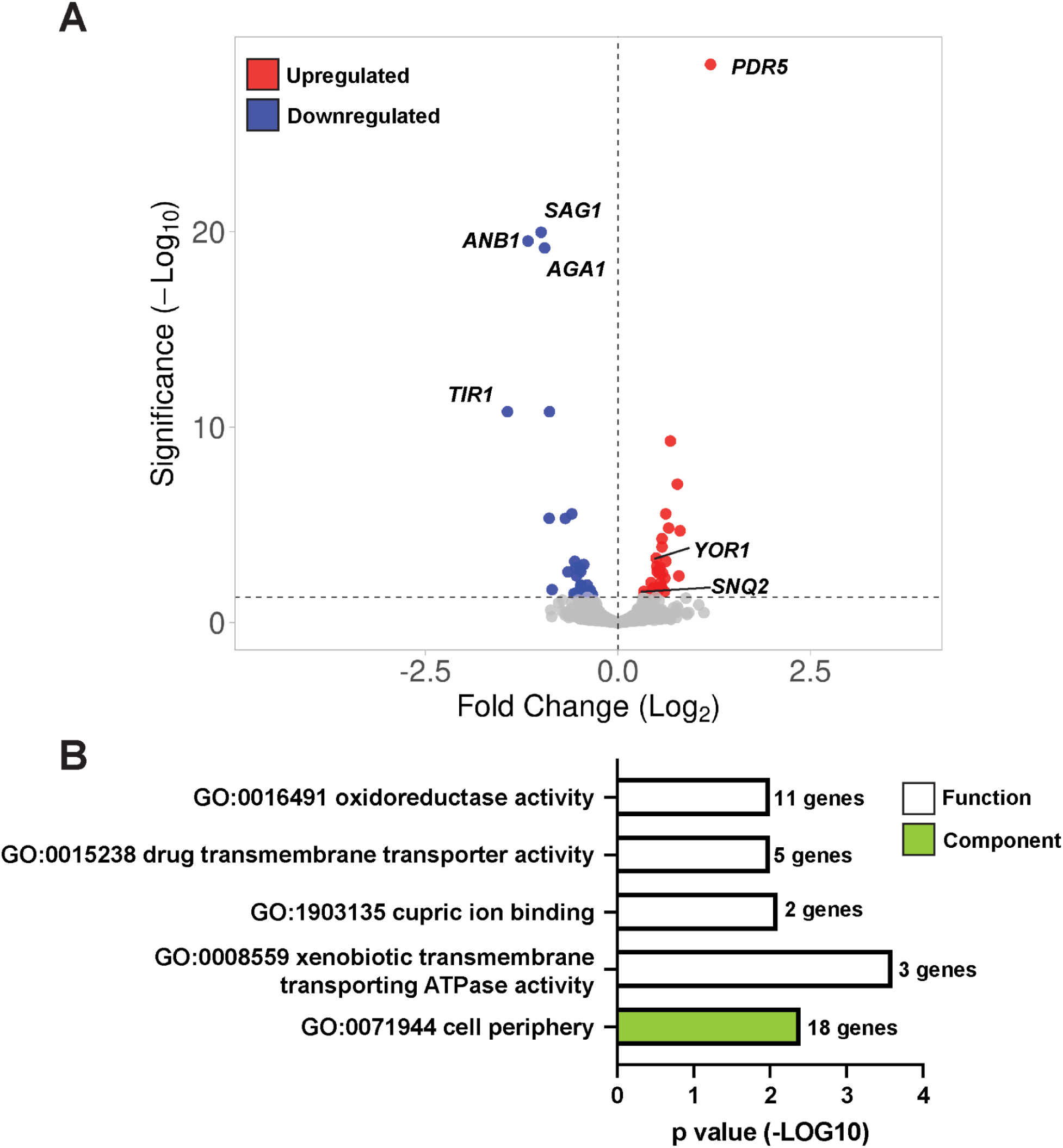
Transcriptional response to TUDCA treatment. **A)** Volcano plot of gene expression determined by RNA-seq from wild-type cells treated with 5 mM TUDCA for 2 h. **B)** Significantly enriched GO biological processes and cellular compartments were determined from the set of significant genes (adjusted *p* < 0.05) in TUDCA-treated cells relative to control.

**Supplemental Figure 2:**
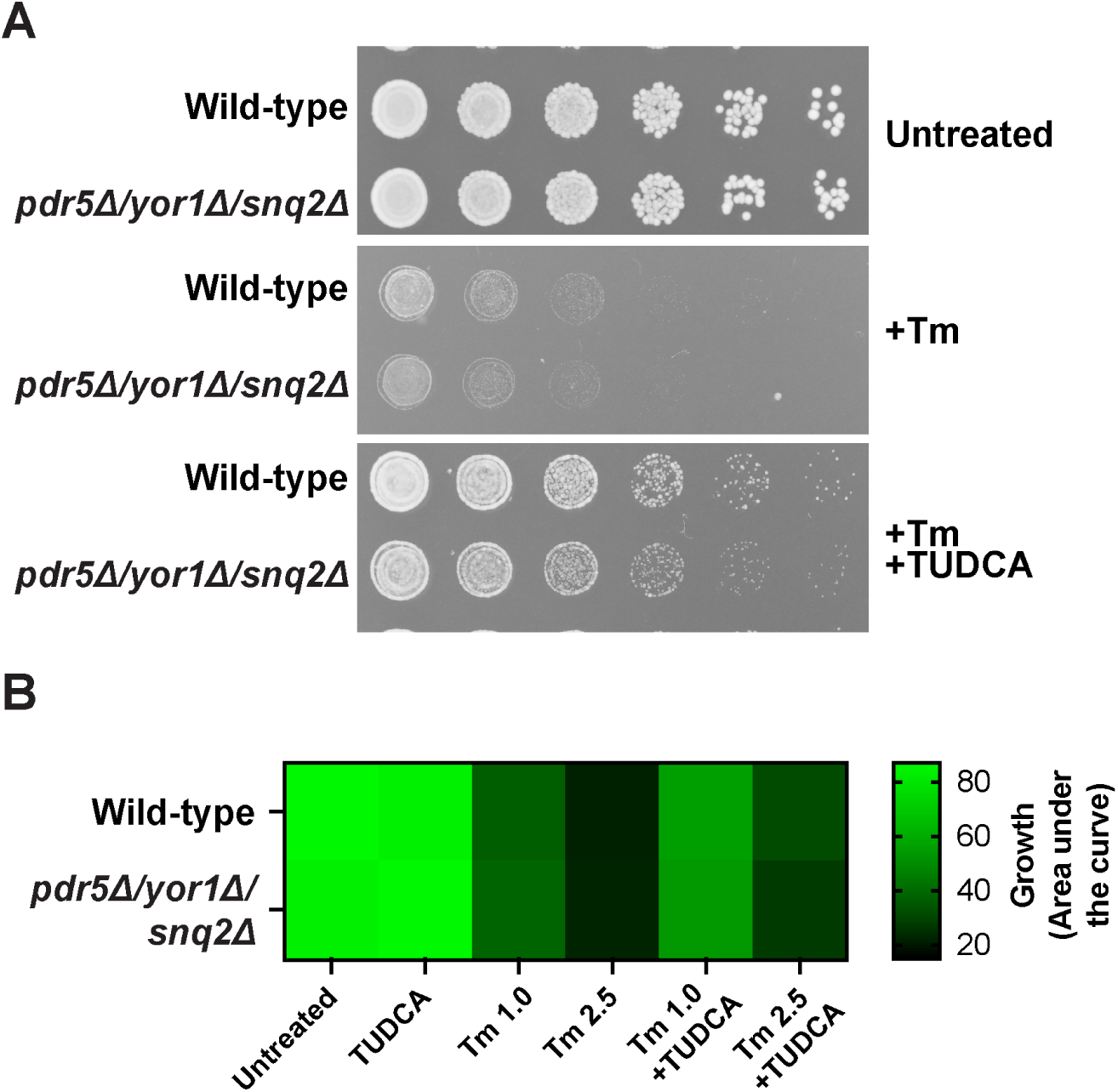
Deletion of *PDR5*, *YOR1* and *SNQ2* multidrug transporters does not impact TUDCA rescue of Tm-induced growth defects. **A)** Wild-type and *pdr5Δ/yor1Δ/snq2Δ* cells were spotted on YPD plates containing 2.5 µg/ml Tm +/− 5 mM TUDCA. Plates were incubated at 30°C for two days before imaging. **B)** Wild-type and *pdr5Δ/yor1Δ/snq2Δ* cells were in liquid YPD media containing 1.0 or 2.5 µg/ml Tm +/− 5 mM TUDCA. Area under the growth curves from liquid cultures over 24 h is presented in a heat map.

**Supplemental Figure 3:**
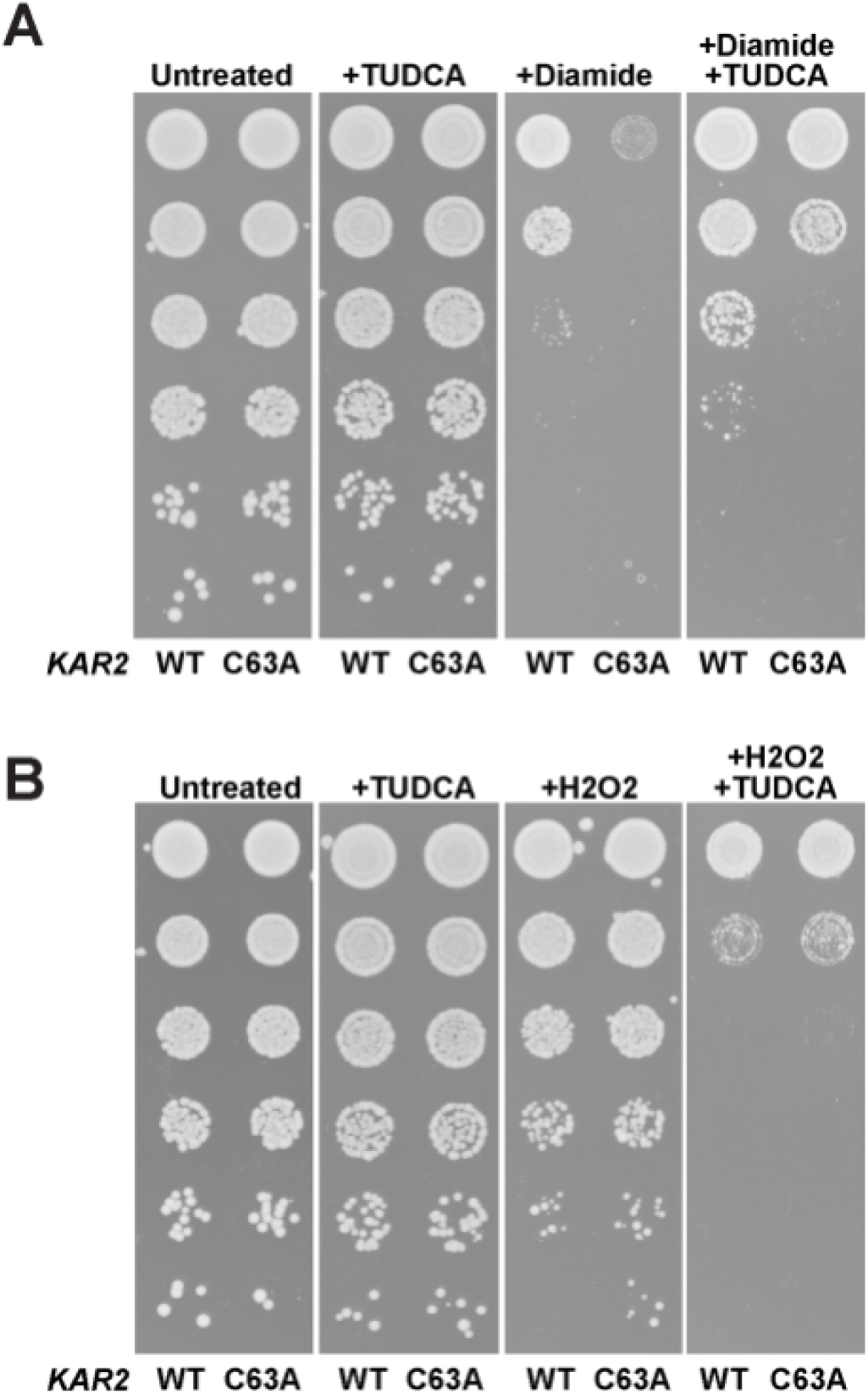
TUDCA improves growth in presence of diamide, but not H_2_O_2_. Cells expressing *KAR2* or *kar2 C63A* were spotted on plates untreated, plus 5 mM TUDCA or containing **A)** 1.25 mM diamide or **B)** 1 mM H_2_O_2_ +/− 5 mM TUDCA. Plates were incubated at 30°C for two days before imaging.

**Supplemental Figure 4:**
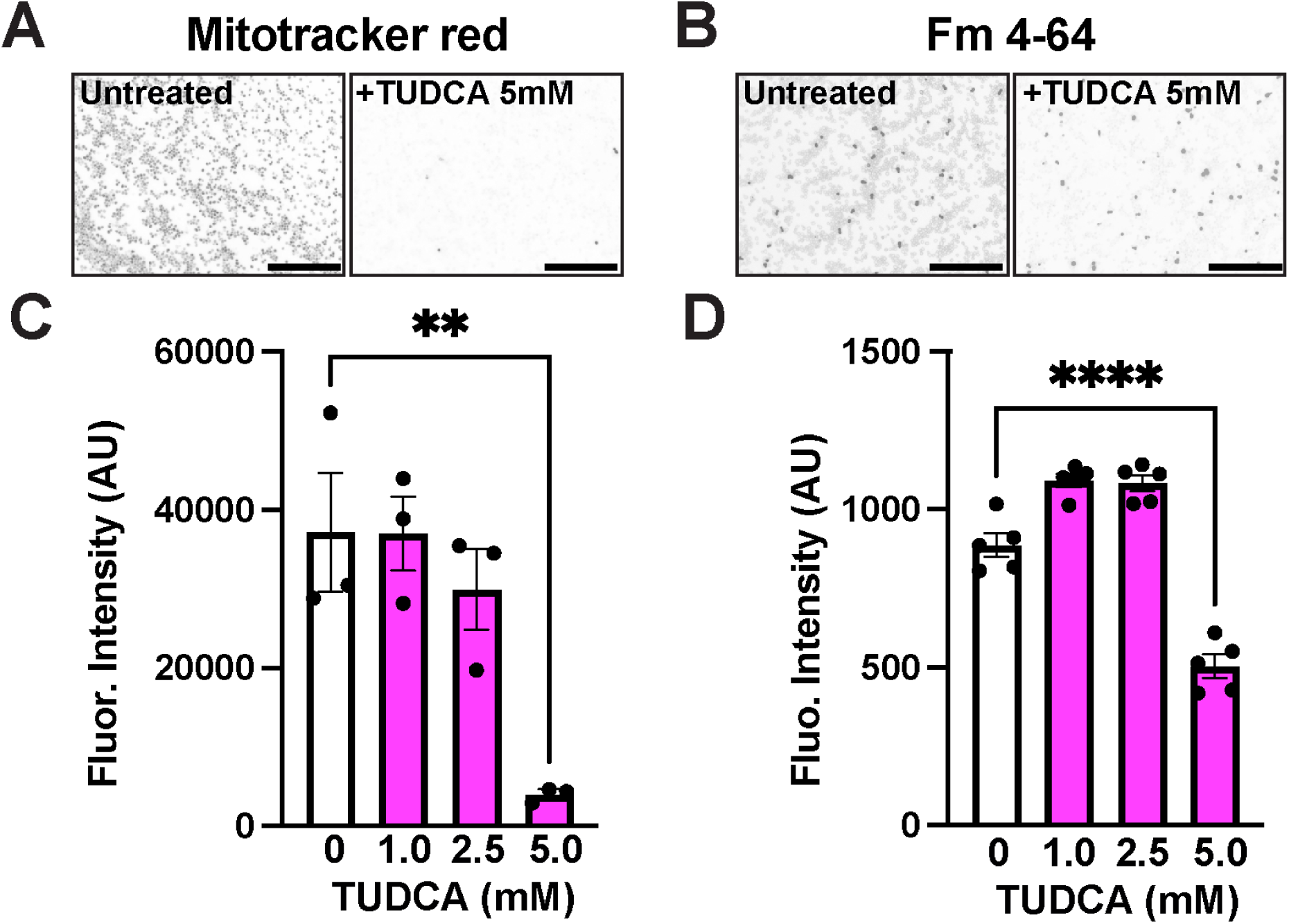
TUDCA impairs staining with some fluorescent dyes. Cells were stained with **(A)** Mitotracker Red (100 nM) or **(B)** FM4-64 (50 µM) in presence or absence or indicated concentration of TUDCA. Cells were imaged immediately using fluorescence microscopy. Representative inverted images are shown. **(C and D)** Fluorescence of both dyes was assessed using flow cytometry. Median fluorescence intensity +/− SEM is shown for 3-5 replicates. **p<0.001 ****p<0.0001. (Anova followed by Tukey’s multi comparison test) Scale bar = 100 µm

**Supplemental Figure 5:**
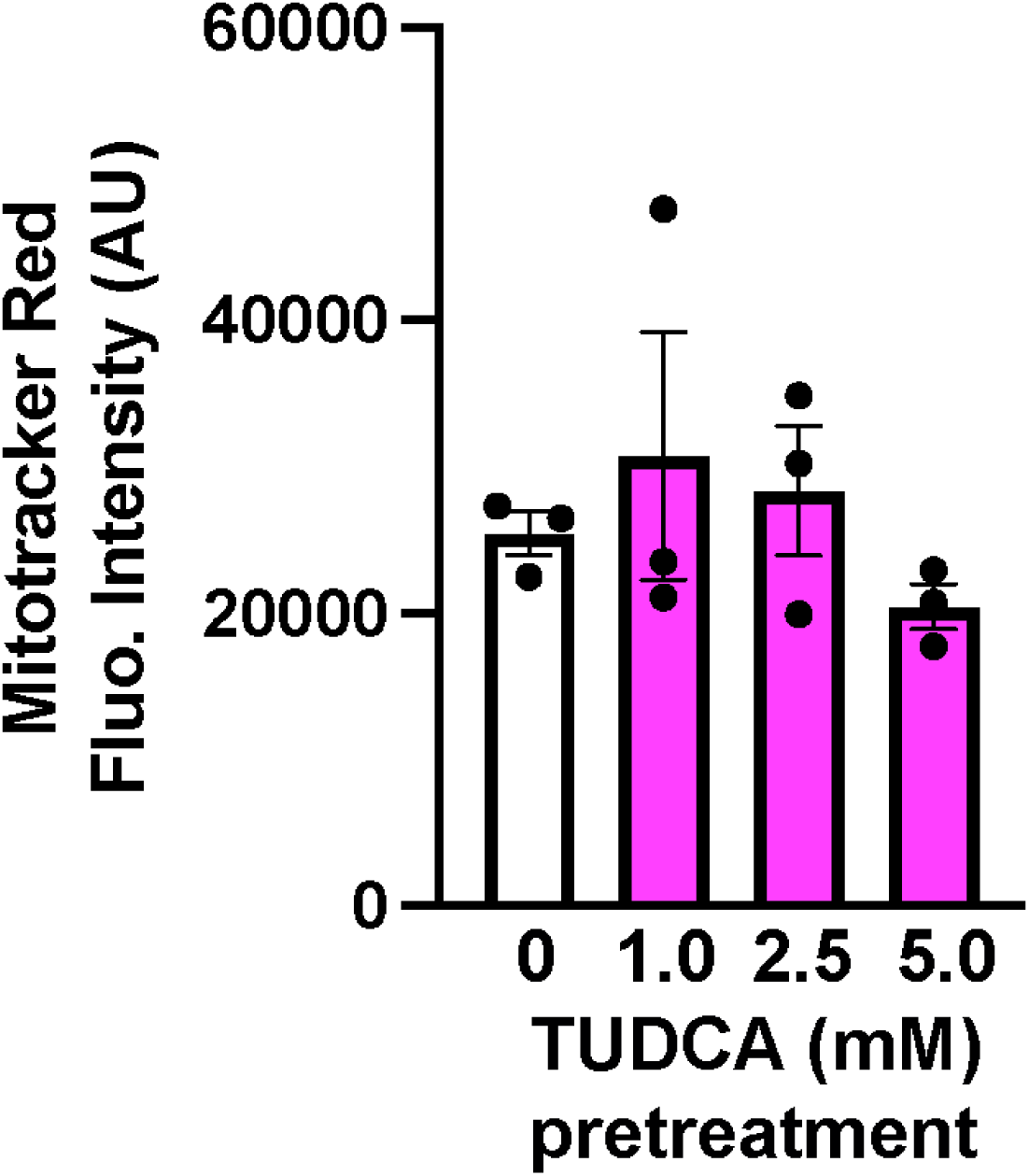
Pretreatment with TUDCA is not sufficient to prevent staining with Mitotracker Red. Cells were pretreated overnight with 1.0, 2.5 or 5.0 mM TUDCA, washed with PBS and stained with Mitotracker Red (100 nM). Fluorescence was assessed using flow cytometry. Median fluorescence intensity +/− SEM is shown for 3 replicates.

**Supplemental Table 1:**
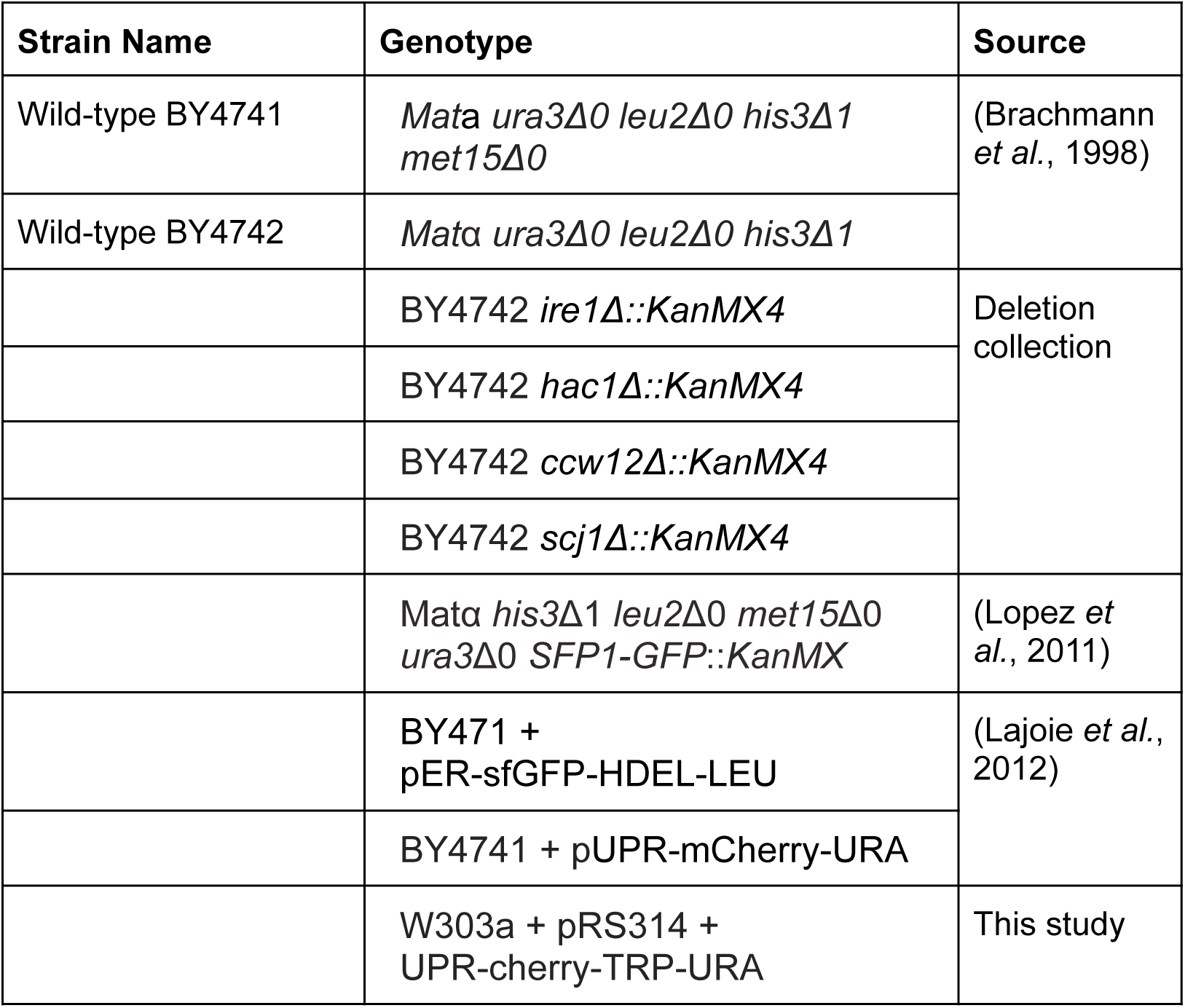
Strains used in this study.

**Supplemental Table 2:** Plasmids used in this study.

| Plasmids | Resistance | Source |
| --- | --- | --- |
| UPR-mCherry<br>Addgene: 20132 | URA | (Merksamer *et al.*, 2008) |
| GPD-yemRFP | LEU | (Albakri *et al.*, 2018) |
| ER-sfGFP-HDEL | LEU | (Lajoie *et al.*, 2012) |
| p415-GAL-CPY | LEU | This study |
| p415-GAL-CPY* |  |  |
| p415-GAL-CPY-ymsfGFP<br>Addgene: 209538 |  |  |
| p415-GAL-CPY*-ymsfGFP<br>Addgene: 209539 |  |  |

**Supplemental Table 3:**
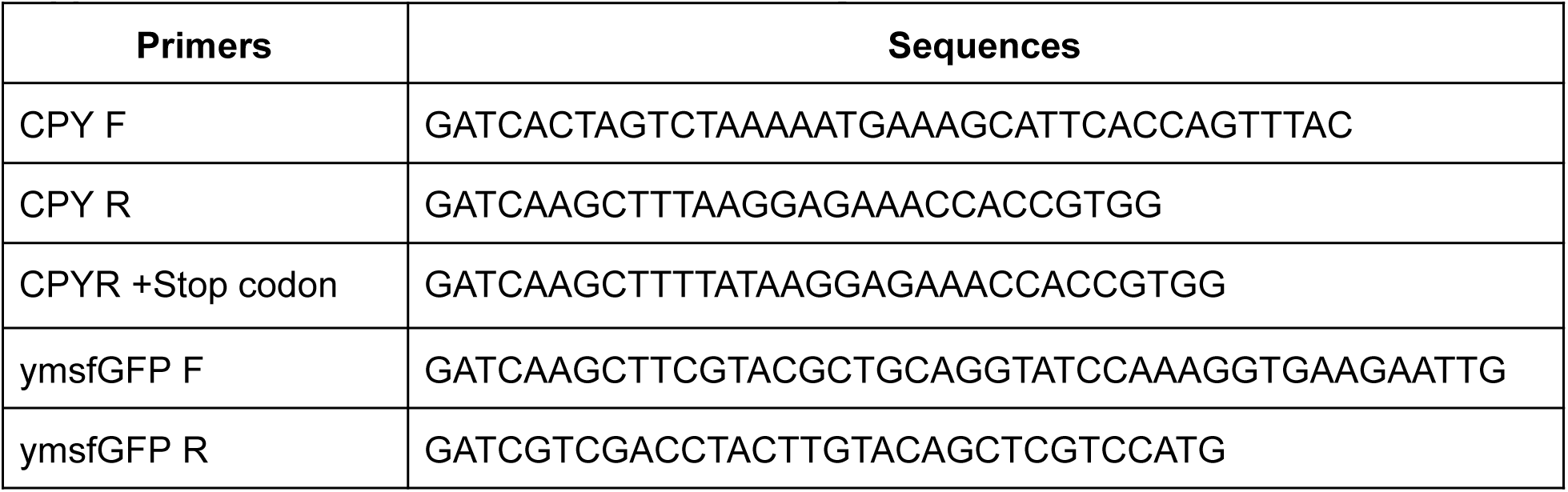
Primers used in this study.

